# Unraveling Hypoxia Tolerance: Transcriptomic and Metabolic Insights from *Lucinoma capensis* in an Oxygen Minimum Zone

**DOI:** 10.1101/2025.08.10.667931

**Authors:** Inna M. Sokolova, Eugene P. Sokolov, Helen Piontkivska, Stefan Timm, Katherine Amorim, Michael L. Zettler

## Abstract

The lucinid clam *Lucinoma capensis* thrives at the oxygen minimum zone margins in the Benguela Upwelling System, where oxygen levels fluctuate dramatically. Understanding its adaptation to such extreme conditions provides key insights into survival strategies under fluctuating oxygen availability. We investigated the transcriptomic and metabolomic responses of *L. capensis* under normoxia, hypoxia, and recovery, focusing on the gills and digestive gland. Our findings highlight distinct organ-specific responses, with the gills showing strong transcriptional changes to oxygen fluctuations, in contrast to the more stable profile observed in the digestive gland. Under hypoxic conditions, the gills exhibited coordinated downregulation of critical metabolic processes, including protein synthesis, transposable element activity, and immune function, suggesting a tightly regulated energy conservation strategy and mechanisms to preserve symbiont stability and genomic integrity. Activation of prokaryotic metabolism in the gills supports the symbionts’ role in host energy acquisition and sulfide detoxification during hypoxia. In contrast, the digestive gland showed minimal transcriptional shifts during anoxia, with upregulation of pathways supporting structural maintenance. Upon reoxygenation, the gills displayed an active and asymmetric recovery, characterized by rapid restoration of protein synthesis and gradual normalization of protein degradation and immune functions. Despite significant transcriptomic changes, the metabolome remained largely stable, reflecting *L. capensis*’s resilience to oxygen fluctuations. However, an overshoot in TCA cycle intermediates and derepression of previously downregulated pathways indicate that reoxygenation involves active metabolic reprogramming, not merely a return to baseline. This study highlights the specialized tissue responses and symbiotic contributions that enable *L. capensis* to thrive in one of the ocean’s most challenging environments.

## 1. Introduction

Since the Great Oxidation Event over 2.2 billion years ago, oxygen (O_2_) has played a key role in shaping life on Earth, driving the evolution of complex organisms and facilitating the development of the biodiversity observed today (Lane, 2002). Oxygen remains critical for the survival of most multicellular organisms due to its central function in cellular bioenergetics (Lane, 2005; Schmidt-Rohr, 2020). However, fluctuations in oxygen availability, such as hypoxia and reoxygenation, impose severe physiological stress on organisms. Hypoxia disrupts oxidative phosphorylation, reducing ATP production and necessitating reliance on less efficient anaerobic metabolism (Hochachka and Mustafa, 1972; Schreurs et al., 2007), while reoxygenation frequently results in excessive production of reactive oxygen species (ROS), causing oxidative stress and cellular damage (Halliwell and Gutteridge, 1999; Hermes-Lima, 2004). These fluctuations affect not only energy metabolism but also broader transcriptional and metabolic networks, requiring adaptive responses that differ across species and habitats (Chen et al., 2024; Connor and Gracey, 2012; Gracey et al., 2008; Lockwood et al., 2015; Yang et al., 2023).

In aquatic environments, oxygen availability is highly variable due to the complex interplay of physical, chemical, and biological processes (Breitburg et al., 2019; Friedrich et al., 2014). Coastal and upwelling systems, in particular, are prone to hypoxia driven by eutrophication, organic matter decomposition, and limited oxygen replenishment (Howarth et al., 2011; Scholz, 2018). Global climate change exacerbates these trends, contributing to widespread deoxygenation by reducing oxygen solubility and enhancing biological respiration, thereby driving biodiversity loss in affected ecosystems (Breitburg et al., 2019; Deutsch et al., 2024). Oxygen minimum zones (OMZs) exemplify such environments, where nutrient-rich upwelling supports high biological productivity, leading to oxygen depletion in subsurface waters (Scholz, 2018).

The Benguela Upwelling System (BUS) is a prominent OMZ characterized by extreme environmental conditions, including periodic hypoxia, hydrogen sulfide accumulation, and high organic matter degradation rates (Amorim and Zettler, 2023; Mohrholz et al., 2008). Seasonal oxygen fluctuations in the BUS range from well-oxygenated conditions in late winter and spring to hypoxic or anoxic states in late summer and autumn, with particularly pronounced variability along the OMZ margins (Zettler et al., 2013). These challenging conditions present significant physiological stresses for benthic marine organisms, yet the margins of the BUS OMZ paradoxically support a highly productive ecosystem dominated by mollusks and polychaetes. Notably, the bivalve mollusks *Lucinoma capensis* and *Lembulus bicuspidatus* achieve exceptional abundances and biomass in these sediments, thriving in hydrogen sulfide-rich muds despite the harsh environmental constraints (Amorim et al., 2021; Eisenbarth and Zettler, 2016; Zettler et al., 2013). While the precise age of the BUS OMZ remains uncertain, evidence suggests that the region’s intense upwelling system has existed for over 3 million years (Mohanty et al., 2024). This prolonged environmental persistence implies that the organisms inhabiting the BUS OMZ may have evolved unique physiological and molecular adaptations to survive and thrive under these extreme and fluctuating conditions. Understanding the molecular physiology of these organisms is critical to elucidating the mechanisms that enable resilience in OMZ ecosystems and contributes to broader insights into the impacts of global deoxygenation on marine biodiversity.

The lucinid clam *L. capensis* exhibits a remarkable capacity to thrive in the hydrogen sulfide-rich sediments of the BUS OMZ, with populations reaching biomass levels of up to 300 g m^−2^ (Amorim et al., 2022). Similar to other lucinids, *L. capensis* forms a symbiotic association with sulfur-oxidizing gammaproteobacteria (*Candidatus Thiodiazotropha*), which detoxify sulfide and supply essential nutrients, enabling the host to inhabit this extreme environment (Amorim et al., 2022; Cary and Felbeck, 1989; Osvatic et al., 2023).While the ecological and symbiotic adaptations of lucinid clams have been extensively studied (Amorim et al., 2022; Cary and Felbeck, 1989; Lim et al., 2019; Osvatic et al., 2023; Yuen et al., 2019), the molecular and physiological mechanisms underlying their responses to severe oxygen fluctuations remain poorly understood.

This study addresses the critical knowledge gap regarding the metabolic physiology of organisms inhabiting OMZs by investigating the transcriptomic and metabolomic responses of *L. capensis* to normoxia (~6.9 ml O_2_ l^−1^), severe hypoxia (~0.11 ml O_2_ l^−1^ for 36 h), and post-hypoxic recovery (1–24 h of normoxia following hypoxia). Two key host tissues were analyzed: the gill, which facilitates gas exchange and hosts sulfur-oxidizing bacterial symbionts, and the digestive gland, which is central to energy storage and metabolism (Amorim et al., 2022; Lim et al., 2019; Yuen et al., 2019). Unlike most previous studies, which primarily focus on the ecological roles and symbiosis of OMZ organisms (Amorim et al., 2022; Cary and Felbeck, 1989; Lim et al., 2019; Osvatic et al., 2023; Yuen et al., 2019), this research provides a rare, in-depth investigation of the host’s molecular and metabolic adaptations to fluctuating oxygen conditions. By examining transcriptomic and metabolomic changes across these conditions, this study provides novel insights into the adaptive strategies that enable *L. capensis* to survive and thrive in the dynamic and extreme environment of the BUS OMZ. The focus on reoxygenation stress—a condition often overlooked but critical for organisms in such habitats—further enhances our understanding of the molecular and physiological mechanisms that mediate environmental resilience of *L. capensis* in OMZ ecosystems.

## 2. Materials and Methods

### 2.1. Chemicals

All chemicals and enzymes were obtained from Biozym Scientific GmbH (Hessisch Oldendorf, Germany), VWR (Darmstadt, Germany), or Fisher Scientific (Schwerte, Germany), and were of analytical grade or higher.

### 2.2. Animal Collection and Maintenance

Adult *L. capensis* (2.1 ± 0.7 cm) were collected at a depth of 150 m by dredging in the central region of the Namibian shelf onboard the FS Meteor during the EVAR Expedition M157 (Fig. 1). A sampling permit was granted by the National Commission on Research, Science and Technology, Namibia (RPIV00812019). During collection, temperature, salinity, and oxygen concentration in the benthic boundary layer were measured using the CTD-system “SBE 911plus” (Seabird-Electronics, USA). In the BUS, *L. capensis* is commonly found in areas of highly variable oxygenation, with dissolved oxygen concentration varying from values < 0.2 ml l^−1^ to <3.4 ml l^−1^ (Amorim & Zettler 2023). Our collection took place during the austral spring (August-September), with oxygen concentrations in *L. capensis* habitats ranging from 0.73 ml O_2_ l^−1^ to 2.4 ml O_2_ l^−1^ during sampling (Amorim et al. 2023). Animals used in this study were collected at approximately 11.7°C, and a dissolved oxygen concentration of 0.73 ml O_2_ l^−1^. All animals used in the experiment were collected in a single dredging from station 12 (23.00°S, 13.87°E, Fig. 1).

**Figure 1.**
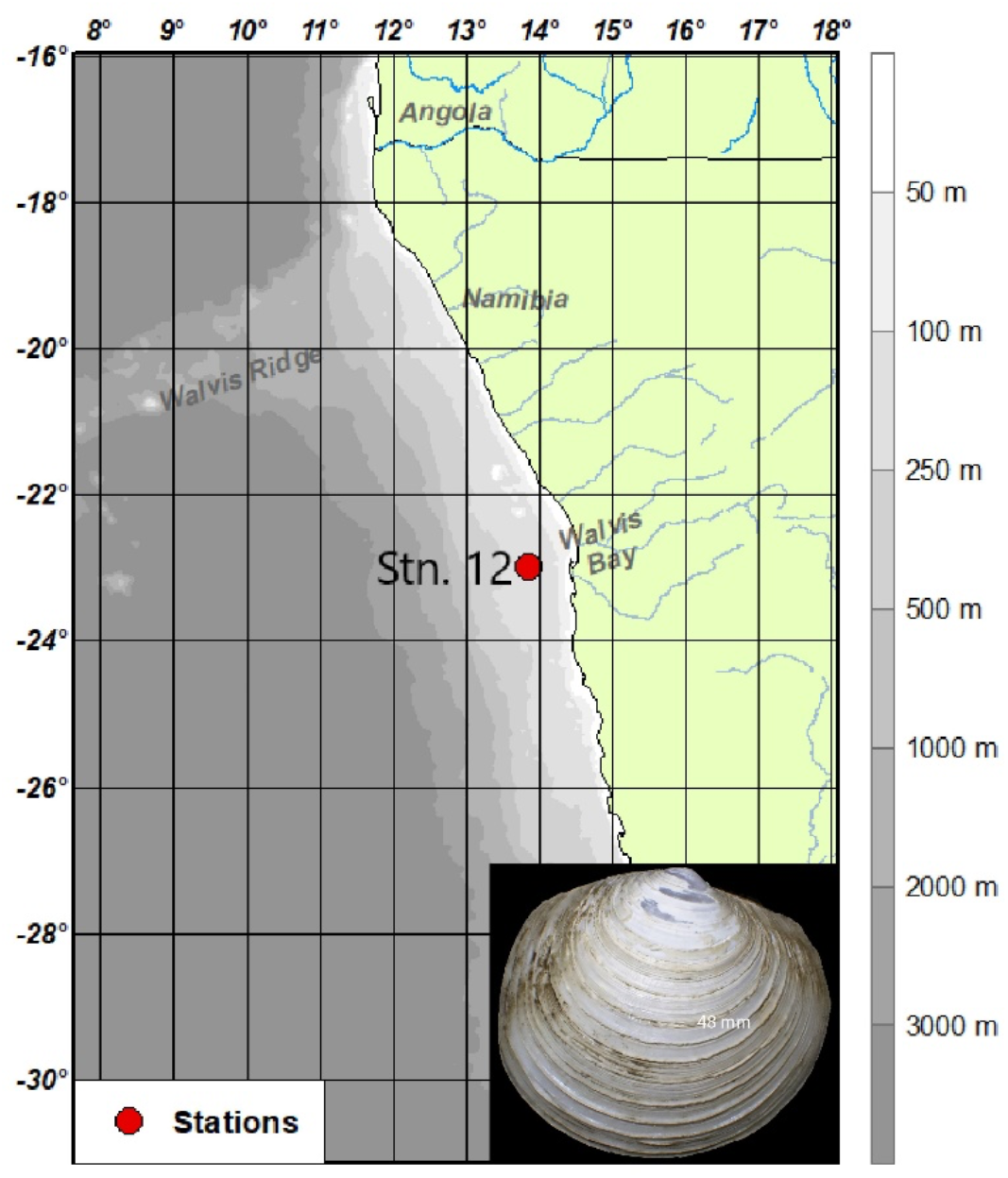
Map of the collection site and ***Lucinoma capensis* photo (insert)**.

### 2.3. Experimental Exposures

Following dredging, 59 individuals of *L. capensis* were randomly distributed into twelve 500 ml glass containers with in situ water at 10°C (1.3 µmol PO_4_ l^−1^; 24.9 µmol NO_3_ l^−1^; 0.26 µmol NO_2_ l^−1^; 0.73 µmol NH_4_ l^−1^; 31.1 µmol SiO_2_). Nutrients concentration in the water was measured in an auto-analyzer (QuaAtro, Seal analytical) using standard calorimetric methods (Grasshoff et al., 2009). Because the bivalves sampled were of highly variable sizes, each bottle received 4-6 individuals, aiming that the sum of the shell lengths of the individuals in each bottle was ca. 10 cm. Twenty hours after sampling, the bottles with the bivalves were assigned to four different conditions (three replicate containers per condition): normoxia (36 h at 6.9 ± 0.24 ml O_2_ l^−1^), severe hypoxia (36 h at 0.11 ± 0.05 ml O_2_ l^−1^), 1 h or 24 h of reoxygenation (36 h of severe hypoxia followed by 1 or 24 h of normoxic recovery). Water in the normoxic treatments was aerated with ambient air. Hypoxia was achieved by continuously bubbling nitrogen through the buffered sea water. O_2_ concentrations were measured using an Optical Oxygen Meter (FireStingO2, Aachen, Germany). All exposures were performed in a temperature-controlled room at 10 °C in the dark. Nutrients concentration were measured in water by the end of exposures only in one bottle of hypoxia condition (2.7 µmol PO_4_ l^−1^; 20.2 µmol NO_3_ l^−1^; 2.2 µmol NO_2_ l^−1^; < 0,5 µmol NH_4_ l^−1^; 34.5 µmol SiO_2_) and two bottles of normoxic condition (0.5 – 1.4 µmol PO_4_ l^−1^; 5.9 – 19.3 µmol NO_3_ l^−1^; 2.6 – 3.8 µmol NO_2_ l^−1^; < 0.5 µmol NH_4_ l^−1^; 32.3 – 33.5 µmol SiO_2_). Except for the lower ammonium levels, other nutrient concentrations were similar to the initial levels in the habitat water.

All bivalves were alive after exposure and were afterwards snap frozen and stored at −80 °C. Frozen tissues (−80°C) were delivered to the Leibniz Institute of the Baltic Sea Research in Rostock and then transported by express service on dry ice to Novogene Sample Receiving Department (Cambridge, U.K.).

### 2.4. Transcriptome Sequencing and Assembly

The RNA extraction, polyA enrichment, mRNA library preparation and next-generation sequencing were carried out by Novogene GmbH (Munich, Germany). Transcriptome sequencing was performed using Illumina NovaSeq 6000 PE150 platform. After adapter trimming, 2 to 4 % of low-quality reads (such as those with over 10% of uncertain nucleotides, or when base quality of less than 5 comprises more that 50 percent of the read) were filtered out, with an average 3% filtered out reads per sample (see Zenodo 10.5281/zenodo.16367847 for the number of reads and other assembly details). The raw sequencing reads were then processed and assembled into transcripts using Trinity 2.6.6 (https://github.com/trinityrnaseq/trinityrnaseq/) (Grabherr et al., 2011). Gene-level counts were computed with CORSET 4.6 (Davidson and Oshlack, 2014) that performs hierarchical clustering of contigs by taking into account shared reads and expression patterns, and selecting the longest transcripts of each cluster as unigenes. (https://github.com/Oshlack/Corset/wiki). For the gill tissue transcriptomic analysis, the sample size (N) was 6 for each experimental condition: normoxia, severe hypoxia, 1-hour recovery, and 24-hour recovery. In the digestive gland, sample sizes were N=2 for the normoxic condition and N=3 for both the severe hypoxia and 24-hour recovery groups. This difference in sample size between the two tissues was due to both biological and technical constraints, including limited tissue availability from some individuals and RNA quality issues that led to the exclusion of samples during library preparation and quality control. Additionally, budget limitations prevented sequencing all samples, so we prioritized gill tissue, which is known to be the most sensitive to hypoxia in bivalves (Lumor et al., 2025; Montúfar-Romero et al., 2024) and thus represents the primary focus of our transcriptomic analysis. The raw sequencing data have been deposited in the NCBI Sequence Read Archive (SRA) under BioProject ID PRJNA1244758, https://www.ncbi.nlm.nih.gov/bioproject/1244758. Supplementary data for transcriptome assembly parameters, including putative functional annotations, Trinity assembly statistics, including N50 and N90, and lengths of transcript and unigene are available in Zenodo <u>10.5281/zenodo.16367847</u>.

### 2.5. Gene Expression Analysis

Gene expression levels were estimated with RSEM (Li and Dewey, 2011) based on transcript abundance using the Corset-filtered Trinity-reconstructed transcriptome as a reference. Read counts were converted into FPKM (Fragments Per Kilobase of transcript sequence per Million base pairs sequenced), an approach that adjusts for both sequencing depth and gene length in fragment counts. Prediction of protein-coding sequences was performed using BLAST (Altschul et al., 1997) against NCBI NR and SwissProt databases (Genbank and UniProt Releases 239 and 2020_04, respectively) for CDS extraction with cut-off e values of 1e-5, and ESTScan with default values (Nagaraj et al., 2007) for unigenes without hits in BLAST.

Differential gene expression analysis was performed with DESeq2 (Love et al., 2014), using read counts as input data. Benjamini–Hochberg correction (Benjamini and Hochberg, 1995) was used to adjust for multiple tests, with the cut-off p-adjusted value of 0.05 and |log2FoldChange| > 0.58 used to identify differentially expressed genes (DEGs) that showed at least 50% change in expression level in either direction.

It is important to note that the library preparation for RNA sequencing was performed using a poly-A enrichment protocol, which selectively captures eukaryotic transcripts and can result in underrepresentation of bacterial mRNAs. Nevertheless, due to the high abundance of symbiotic and associated bacteria in gill tissues, a substantial number of prokaryotic transcripts were detected (Supplementary Table 3). In contrast, the digestive gland—lacking intracellular symbionts and containing only a typical extracellular microbiome—showed no detectable prokaryotic differential expression. However, without targeted microbial metatranscriptomics or complementary metagenomic sequencing, we cannot confidently assign these transcripts to specific microbial taxa. As such, our interpretation of prokaryotic gene expression is limited to broad functional trends rather than taxon-specific insights.

Gene functional enrichment analysis was then performed to identify biological functions or pathways significantly associated with DEGs using biological ontologies, Gene Ontology (GO) and Kyoto Encyclopedia of Genes and Genomes (KEGG), using GOseq (https://bioconductor.org/packages/release/bioc/html/goseq.html) (Young et al., 2010) and KOBAS (Wu et al., 2006), respectively, with corrected p values < 0.05. For a more in-depth insight into the functional transcriptomic response to hypoxia-reoxygenation, we manually curated the list of DEGs by assigning all annotated DEGs to 64 functional categories for eukaryotic transcripts and 41 categories for prokaryotic transcripts (Supplementary Table 1). The transcripts annotated as genes with unknown functions were omitted. Each DEG was manually reviewed and assigned to a functional category based on its annotated function, supported by homology searches against the NCBI nucleotide database, the SwissProt protein database, and the KEGG Orthology (KO) database (see Zenodo 10.5281/zenodo.16367847 for details of manual annotation). The likely origin of each gene—prokaryotic or eukaryotic—was determined through sequence homology comparisons with relevant reference databases. To ensure non-redundancy, genes sharing identical NCBI or SwissProt accession numbers were flagged, reviewed, and, where appropriate, merged or removed. This process yielded a final non-redundant list of DEGs. Functional categories were then quantified by counting the number of genes assigned to each category. Finally, a summary table of DEG counts per merged functional category was generated using standard Excel functions, including lookups and conditional counting.

#### Reactome analysis

Since Reactome overrepresentation algorithm makes pathway predictions based on the human genomic database (Milacic et al., 2023), we converted non-human gene IDs to the Ensembl IDs of respective human homologs using BLASTP search against protein sequences of GRCh38 human genome assembly (https://ftp.ensembl.org/pub/release-110/fasta/homo_sapiens/pep/). We used relatively relaxed cut-off e value of 1e-5 to capture even distant homologs. Only eukaryotic transcripts of *L. capensis* were included in search of homologs.

Once we identified respective human homologs, we applied two cut-off thresholds for Reactome pathways overrepresentation interpretation: a more conservative FDR < 0.05 and a more relaxed FDR < 0.1. Setting the FDR at 0.1, instead of the traditional 0.05, increases sensitivity, allowing us to capture more potentially relevant metabolic pathways. In exploratory studies or large-scale analyses with numerous tests, such as ours, a less stringent FDR helps detect subtle but meaningful patterns, minimizing the risk of overlooking true positives (Benjamini and Hochberg, 1995). This approach is particularly valuable when the focus is on identifying promising leads for further investigation, rather than confirming established hypotheses. Reactome analyses were performed using version 90 (released September 2024).

We would like to acknowledge that using Reactome, which is focused on human data for pathway annotations, has limitations when applied to non-model organisms. These limitations include potential discrepancies in physiological processes—such as the presence or absence of an adaptive immune response—between mammals and bivalves. However, Reactome offers a broad and detailed set of insights, manually curated by domain experts, into both normal and disease-related pathways (Milacic et al., 2024; Rothfels et al., 2023). As such, it provides valuable opportunities for identifying evolutionarily conserved aspects of fundamental molecular and cellular processes, such as those involved in the essential response to hypoxia (O’Farrell, 2001; Semenza, 2004; Webster, 2003).

### 2.6. Metabolite Analysis

Targeted metabolome analysis was conducted as described elsewhere (Bruhns et al., 2023). Briefly, metabolites were extracted from the gill (mean tissue mass: 37.0±3.5 mg, range 3.5-98.9 mg) and the digestive gland (mean tissue mass: 35.4±5.4 mg, range 2-95.5 mg) tissue, homogenized in 1 ml of ice-cold 80% ethanol containing 1 μg ml^−1^ of 2-(N-morpholino)ethanesulfonic acid (MES) as an internal standard. Each sample was derived from individual clams. The homogenates were centrifuged at 13,000 ×g for 10 min at 4°C to remove debris. The resulting supernatants were freeze-dried using a cold trap (Unicryo MC 2 L, UniEquip, Germany) and stored at −80°C for future analysis. Prior to measurement, extracts were reconstituted in LC-MS-grade water (ROTISOLV, Carl Roth) and passed through sterile 0.2 μm filters (Omnifix-F, Braun, Germany). Metabolite measurements were conducted on a Shimadzu LCMS-8050 high-performance liquid chromatography-mass spectrometer. Metabolites were identified and quantified using the built-in LC-MS/MS software package for primary metabolites (Version 2, Shimadzu, P/N 225–24,862-92) and LabSolutions software (Shimadzu). Calibration was performed using metabolite-specific standards from Merck, with normalization based on the internal standard (2-(N-morpholino)ethanesulfonic acid). In total, 35 metabolites - including amino acids, intermediates of the tricarboxylic acid (TCA) cycle and urea cycle, adenosine monophosphate (AMP), S-adenosylmethionine, carnitine, nicotinamide adenine dinucleotide (NAD) and γ-aminobutyric acid (GABA) - were measured. Concentrations of iso(citrate), aconitate and fumarate were below the detection limits of the method. In total, 31 and 34 samples were processed for the gill and the digestive gland, respectively. For the metabolomic analysis, sample sizes (N) in gill tissue were as follows: 8 for normoxia, 11 for severe hypoxia, 7 for 1-hour recovery, and 5 for 24-hour recovery. In the digestive gland, sample sizes were N=8 for normoxia, 11 for severe hypoxia, 8 for 1-hour recovery, and 7 for 24-hour recovery.

Data distribution and variance were assessed using normal probability plots and Bartlett’s test, which is sensitive to deviations from normality and homoscedasticity. For succinate levels in gills, zero values were replaced by 1/2 of the lowest positive value considered a limit of detection (4.21 ng mg^−1^). Data for all metabolites except AMP met the criteria for normality and homogeneity of variances and were analyzed via one-way ANOVA (oxygen condition as a fixed factor), followed by Tukey’s Honest Significant Differences test for post-hoc comparisons. For AMP concentrations, non-parametric Kruskal-Wallis test was applied, with Dunn’s test used for multiple comparisons. The ANOVA and Kruskal-Wallis analyses were conducted using GraphPad Prism ver. 7.05 (GraphPad Software Inc., Boston, MA, USA).

Multivariate analyses of the metabolite profiles, including sparse partial least squares – discriminant analysis (sPLS-DA), correlation analysis and clustering, were performed on data that were autoscaled (mean-centered and divided by the standard deviation of each variable) to minimize the effects of differences in absolute metabolite concentrations as implemented in MetaboAnalyst 6.0 (Pang et al., 2024). The sPLS-DA algorithm is well-suited for datasets where the number of variables (metabolites) significantly exceeds the number of samples, as it helps reduce the variable set to generate robust and easily interpretable models (Lê Cao et al., 2011). In our model, we specified a maximum of five principal components, with 10 variables selected per component. The effects of oxygen condition on tissue-specific metabolism were assessed using pathway enrichment analysis in MetaboAnalyst 6.0 (Pang et al., 2024). This approach identifies subtle yet consistent changes in metabolites within the same biological pathway. The *Drosophila melanogaster* metabolic reference library and the Globaltest method were used for pathway enrichment, with relative betweenness centrality as the measure of node importance. Compounds were autoscaled by their means and standard deviations, and pathways with a false discovery rate (FDR) <0.05 and a pathway impact >0 were considered significant for further analysis. All analyses were conducted using GraphPad Prism v. 7.02 (GraphPad Software Inc., La Jolla, CA, USA) and Metaboanalyst 6.0 (https://www.metaboanalyst.ca/) (Pang et al., 2024), with differences considered significant at p<0.05. The metabolomics data are available via Zenodo, <u>10.5281/zenodo.15268760</u>.

## 3. Results

### Transcriptome overview

A comparison of the whole transcriptome data across all replicates revealed a substantial number of co-expressed transcripts among different exposure conditions in both studied tissues. Specifically, 68,267 transcripts in the gills and 55,981 transcripts in the digestive gland were shared across all treatments (Fig. 2). In the gills, the highest number of uniquely expressed transcripts was observed after 1 h of reoxygenation (10,491), followed by the normoxic controls (10,088), 24 h of reoxygenation (9,789), and the lowest during hypoxia (8,121). In the digestive gland, the greatest number of uniquely expressed transcripts occurred in the 24-h recovery group (19,936), followed by the hypoxia group (12,344) and the normoxic control (10,822).

**Figure 2.**
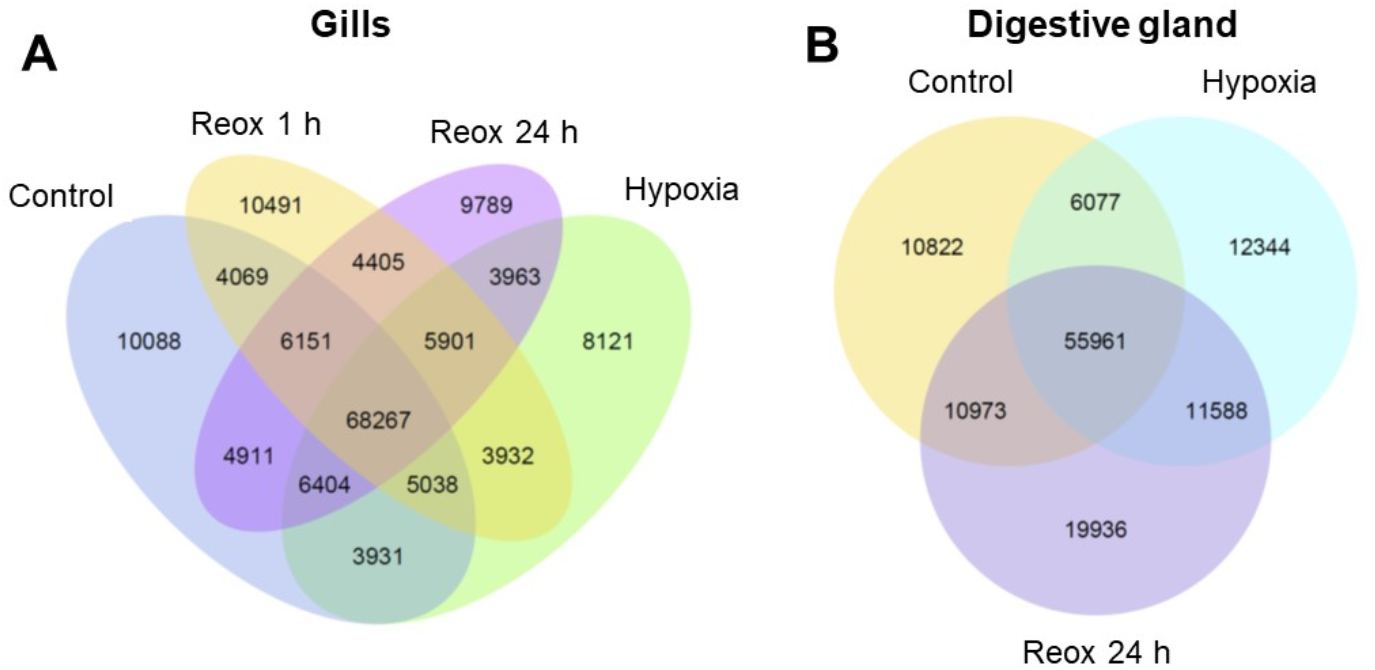
Venn diagram of the gill (A) and the digestive gland (B) transcriptome of *L. capensis*. Reox – reoxygenation. Numbers inside the respective sections indicate the number of uniquely expressed or co-expressed transcripts.

### Differential gene expression: Summary

Using the selected cut-off criteria (p_adj_ < 0.05, |log2FoldChange| > 0.58), the highest number of DEGs, totaling 9,006, was found when comparing the gill transcriptome of *L. capensis* under normoxic conditions to that of the clams exposed to hypoxia. Of these DEGs, 1,500 were upregulated, and 7,506 were downregulated (Fig. 3A). After 1 h of post-hypoxia recovery, the gill transcriptome remained significantly altered compared to the normoxic state, showing 6,501 DEGs, with 1,436 upregulated and 5,065 downregulated transcripts. Following 24 h of reoxygenation, 4,214 gill transcripts were differentially expressed relative to the normoxic controls, with 1,942 upregulated and 2,272 downregulated.

**Figure 3.**
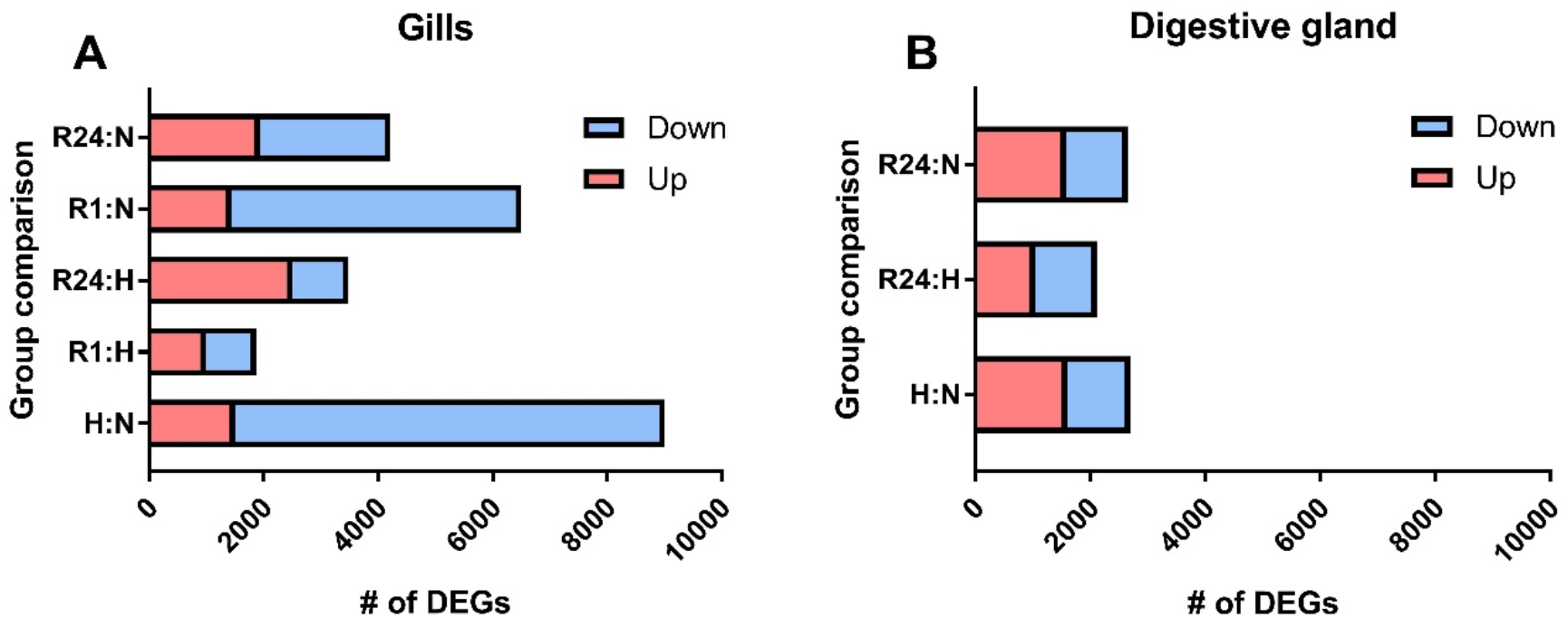
DEG counts in the gills (A) and the digestive gland (B) of *L. capensis*. Experimental conditions: H – hypoxia, N – normoxic control, R1 – 1 h of post-hypoxia recovery, R24 – 24 h of post-hypoxia recovery. The DEGs are shown for all pairwise comparisons of oxygen regimes within the two studied tissues.

In the digestive gland, 2,697 genes (1,597 upregulated and 1,100 downregulated) were differentially expressed after hypoxia, and 2,653 genes (1,578 upregulated and 1,075 downregulated) were observed after 24 h of recovery compared to the normoxic controls (Fig. 3B).

Many of the identified DEGs were annotated as genes with unknown functions and were therefore excluded from further analysis. The subsequent analysis focused on DEGs with putative functions that could be identified through homology with known proteins from other species.

### Differential gene expression: Gills

In the hypoxic gills of *L. capensis*, 819 eukaryotic host transcripts were downregulated, while 236 were upregulated compared to normoxic controls (Supplementary Table 2A, B). Among the downregulated transcripts, processes contributing more than 5% to the DEG pool included protein synthesis and breakdown, mobile genetic elements, immunity, and signaling. Conversely, the upregulated transcripts with greater than 5% representation were associated with proteolysis, transcription regulation, immunity, cytoskeleton organization, carbohydrate metabolism, DNA maintenance, extracellular matrix, and protein synthesis. A marked imbalance was observed among functional groups, with protein synthesis showing a 12.5-fold excess of downregulated over upregulated DEGs, followed by mobile genetic elements (8.3-fold), signaling (5.9-fold), and immunity (3.1-fold).

During the recovery phase after 1 h of post-hypoxia reoxygenation, 409 eukaryotic host transcripts were downregulated, and 289 were upregulated (Supplementary Table 2C, D). The primary functional categories (>5% of DEGs) among the downregulated transcripts were proteolysis, mobile genetic elements, immunity, transcription regulation and protein synthesis. The upregulated transcripts were mostly associated with proteolysis, extracellular matrix, mobile genetic elements, immunity, transcription regulation, and signaling. Apoptosis exhibited the most pronounced downregulation, with an over 8-fold excess of downregulated compared to upregulated transcripts, whereas the extracellular matrix showed a 3.2-fold excess of upregulated transcripts.

Following 24 h of reoxygenation, 261 eukaryotic host transcripts were downregulated, while 328 were upregulated (Supplementary Table 2E, F). Among the downregulated DEGs, the most prominent (>5%) functional groups included immunity, proteolysis, protein synthesis, and mobile genetic elements. The upregulated DEGs were predominantly associated with proteolysis, transcription regulation, DNA maintenance, mobile genetic elements, and signaling. The most substantial upregulation was observed in DNA maintenance, with a 3.8-fold excess of upregulated over downregulated transcripts.

Due to the use of polyA enrichment during library preparation, which selectively captures eukaryotic transcripts, bacterial transcripts lacking polyadenylation may be substantially underrepresented in our dataset. As such, the prokaryotic DEGs reported here likely reflect only a partial view of the symbiont’s transcriptional response. Nevertheless, for prokaryotic symbiont transcripts detected in the hypoxic gills, 40 were downregulated and 139 were upregulated compared to normoxic conditions (Supplementary Table 3A, B). Downregulated transcripts (>5% of DEG) were primarily associated with antibiotic and amino acid metabolism, and proteolysis. Among the upregulated prokaryotic DEGs, the most significant functional groups included sulfur metabolism, CO_2_ fixation, electron transport system (ETS), protein synthesis, and amino acid metabolism. Notably, the greatest upregulation compared to downregulation was observed in ETS (16-fold), protein synthesis (15-fold), and CO_2_ fixation (9-fold).

After 1 h of reoxygenation in the gills of *L. capensis*, 88 prokaryotic symbiont transcripts were downregulated, while 52 were upregulated (Supplementary Table 3C, D). The downregulated transcripts were primarily involved in amino acid metabolism, protein synthesis, and mobile genetic elements. In contrast, upregulated transcripts were predominantly associated with CO_2_ fixation, nitrogen and sulfur metabolism, transcription regulation, the tricarboxylic acid cycle, and amino acid metabolism.

Following 24 h of reoxygenation, 18 prokaryotic symbiont transcripts were downregulated, and 84 were upregulated (Supplementary Table 3E, F). No dominant functional groups were identified among the downregulated DEGs, as most consisted of only 1–2 transcripts. The upregulated transcripts were enriched in protein synthesis, CO_2_ fixation, transcriptional regulation, sulfur metabolism, ETS, and chaperones, along with amino acid metabolism.

### Differential gene expression: Digestive gland

After hypoxia, we identified and functionally annotated 167 downregulated and 382 upregulated eukaryotic DEGs in the digestive gland of *L. capensis* compared to the normoxic control (Supplementary Table 4A, B). Among the downregulated DEGs, the most prominent functional categories (>5% of DEGs) were related to protein synthesis, mobile genetic elements and proteolysis. The upregulated DEGs were primarily associated with immunity, proteolysis and extracellular matrix. The largest excess of upregulated over downregulated transcripts was observed in the functional categories related to immunity (8.9-fold) and proteolysis (3.5-fold).

In the digestive gland of *L. capensis* following 24 h of reoxygenation, 144 transcripts were downregulated, while 384 were upregulated compared to normoxic conditions (Supplementary Table 4C, D). The most prominent downregulated functional categories included protein synthesis, immunity, and mobile genetic elements. In contrast, upregulated transcripts were predominantly associated with protein synthesis, immunity, proteolysis, and neural function. Among these, the functional category related to proteolysis exhibited the most pronounced imbalance, with a 3.6-fold excess of upregulated over downregulated transcripts. No prokaryotic DEGs were identified in the digestive gland of *L. capensis*. Given the use of polyA enrichment, which biases against non-polyadenylated bacterial transcripts, the absence of detectable prokaryotic DEGs in the digestive gland of *L. capensis* should be interpreted with caution. However, this result is consistent with the known lack of endosymbionts in this tissue (Amorim et al., 2022).

### GO and KEGG pathway enrichment analysis

Exposure to hypoxia in the gills of *L. capensis* resulted in the significant downregulation of numerous GO pathways (padj < 0.05), particularly those linked to protein metabolism and cellular biosynthesis (Fig. 4A). The regulation of mobile genetic elements (transposition) was also notably enriched among downregulated DEGs. Conversely, only three GO pathways—associated with endosymbiont metabolism (plastid, photosynthesis, and oxidoreductase activity)—were significantly enriched among the upregulated genes (padj < 0.05) (Fig. 4B). The DEGs associated with plastid and photosynthesis GO pathways included several subunits of ribulose-1,5-bisphosphate carboxylase/oxygenase (RuBisCO), a key enzyme in in carbon fixation via the Calvin–Benson–Bassham cycle.

**Figure 4.**
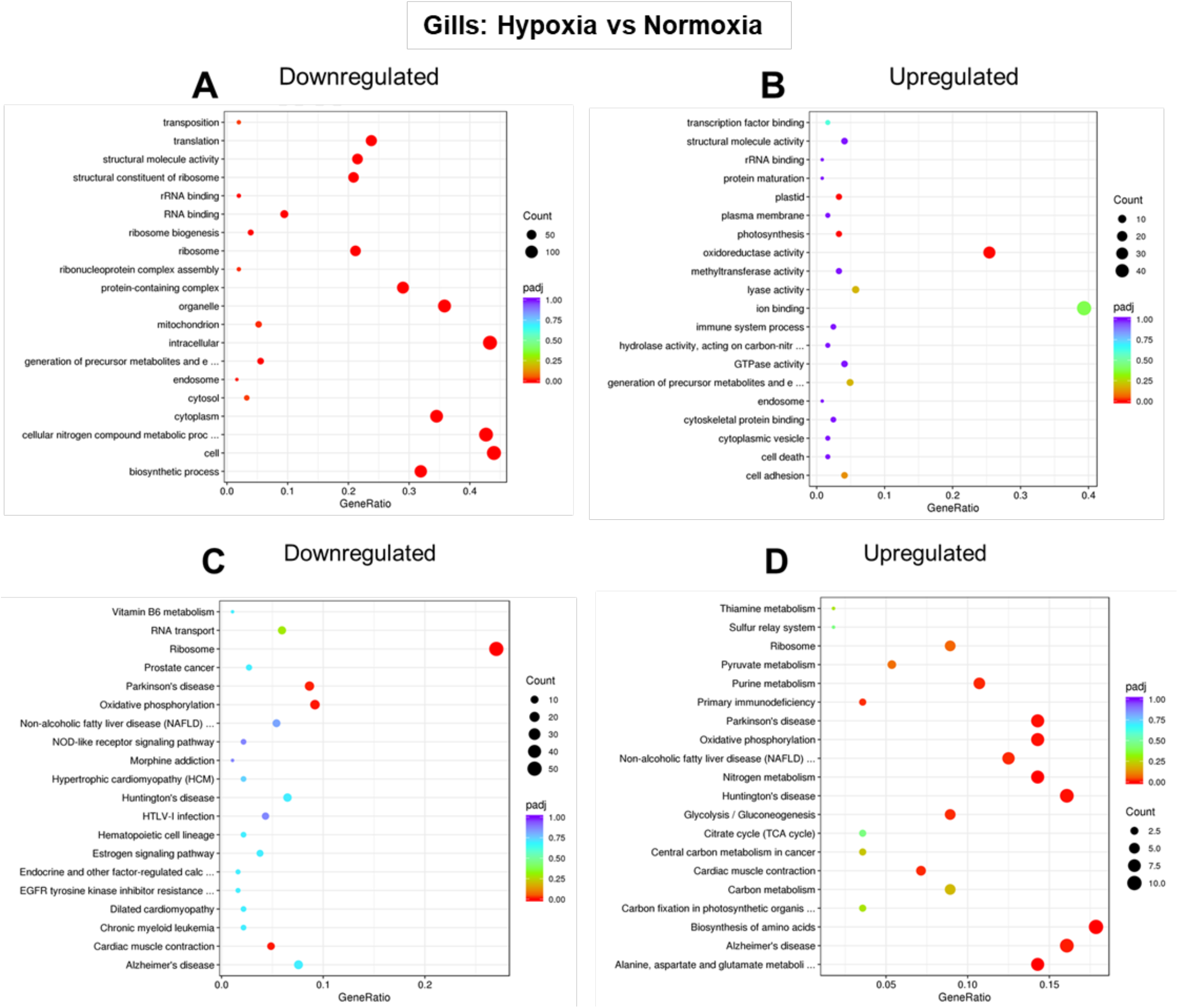
GO (A, B) and KEGG (C,D) pathway enrichment in the gills of hypoxia *L. capensis* relative to the normoxic control. Color indicates the significance (p_adj_) and size of the symbols – the number of DEGs found in the respective pathway. X axis shows Gene Ratio for each pathway, depicting the ratio of differentially expressed genes to all genes for this GO term or KEGG pathway.

KEGG enrichment analysis highlighted the downregulation of pathways related to ribosome function, oxidative phosphorylation, cardiac muscle contraction, and Parkinson’s disease (padj < 0.05) (Fig. 4C). In contrast, upregulated KEGG pathways included various metabolic processes (e.g., nitrogen metabolism, biosynthesis of amino acids, and glycolysis/gluconeogenesis), oxidative phosphorylation, cardiac muscle contraction, and several disease-related KEGG pathways (padj < 0.05) (Fig. 4D). Since KEGG annotations are based on human biology, disease-related pathways up- or downregulated in clams do not indicate human-like pathologies, but rather reflect changes in DEGs associated with evolutionary conserved processes such as mitochondrial function, protein degradation, metabolism, and apoptosis (see Zenodo dataset 10.5281/zenodo.16367847 for DEG annotation).

After 1 h of reoxygenation, no significantly downregulated GO nor KEGG pathways were detected in the gills of *L. capensis* compared to the normoxic state (Fig. 5A, C). However, two GO pathways associated with endosymbiont metabolism (plastid, and photosynthesis both involving RuBisCO subunits) were significantly enriched among upregulated genes (padj < 0.05) (Fig. 5B). After 24 h of reoxygenation, the transposition GO pathway was significantly downregulated (p_adj_<0.05) relative to the normoxic state in the gills of *L. capensis* (Fig. 6A). Two GO pathways (chromosome and histone binding) were significantly enriched (p_adj_<0.05) among the upregulated genes in *L. capensis* gills after 24 h of recovery (Fig. 6B). Across both recovery time points, no significantly downregulated or upregulated KEGG pathways were found in the gills relative to normoxic conditions (Fig. 5C, D, Fig. 6C, D, and Supplementary Figure 2).

**Figure 5.**
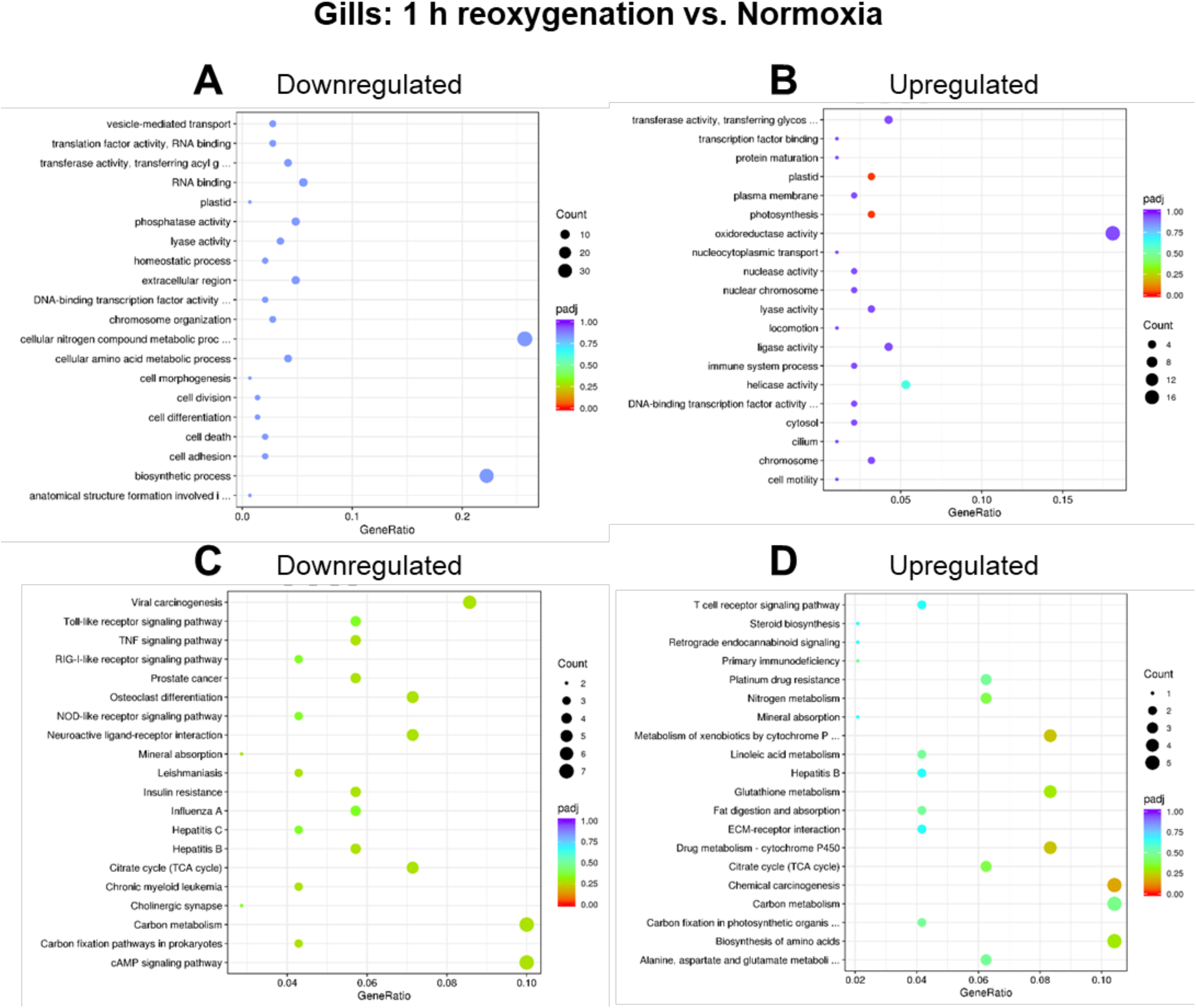
GO (A, B) and KEGG (C,D) pathway enrichment in the gills of *L. capensis* after 1 h of reoxygenation relative to the normoxic control. Color indicates the significance (p_adj_) and size of the symbols – the number of DEGs found in the respective pathway. X axis shows Gene Ratio for each pathway, depicting the ratio of differentially expressed genes to all genes for this GO term or KEGG pathway.

**Figure 6.**
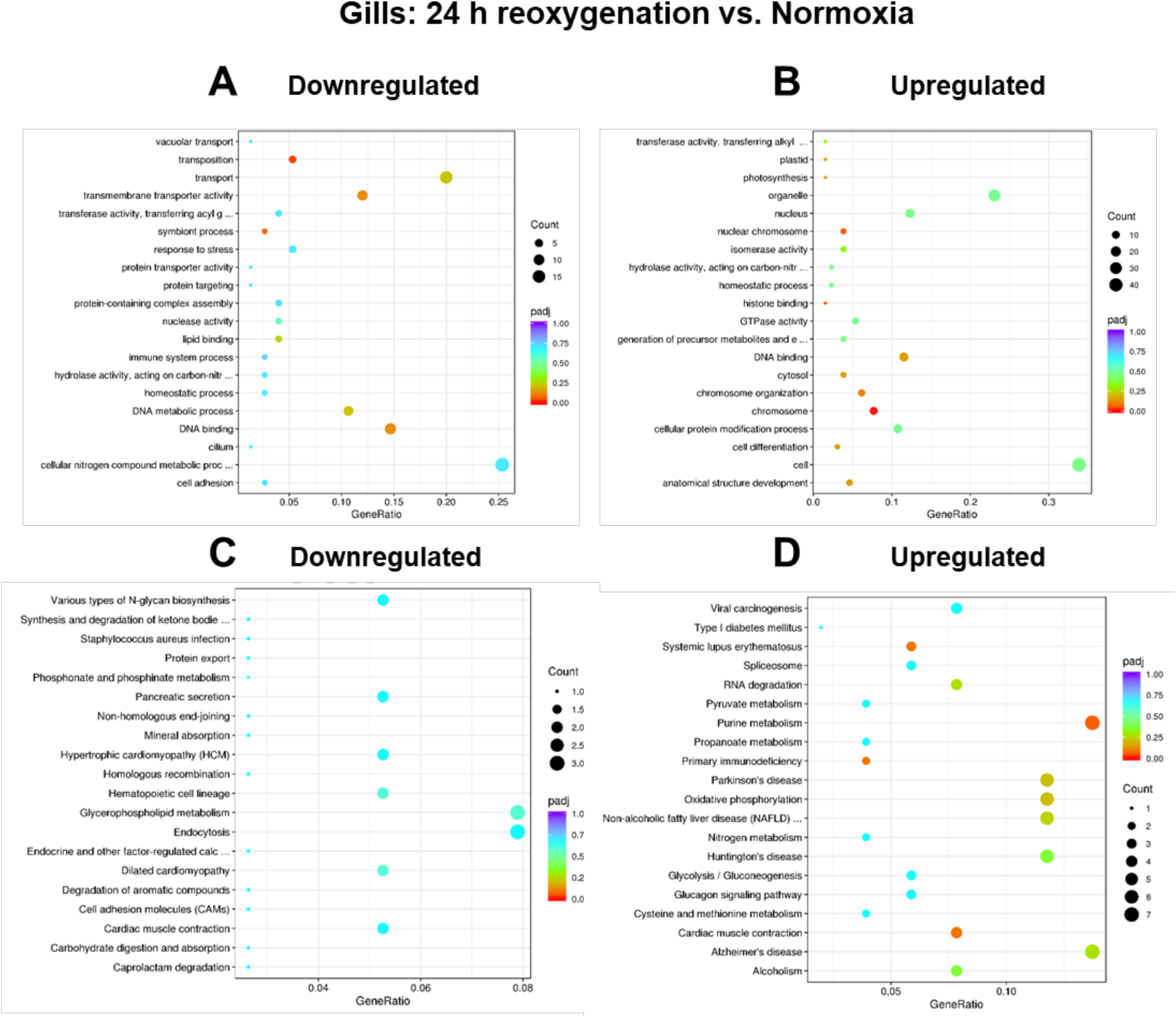
GO (A, B) and KEGG (C,D) pathway enrichment in the gills of *L. capensis* after 24 h of reoxygenation relative to the normoxic control. Color indicates the significance (p_adj_) and size of the symbols – the number of DEGs found in the respective pathway. X axis shows Gene Ratio for each pathway, depicting the ratio of differentially expressed genes to all genes for this GO term or KEGG pathway.

In the digestive gland tissue, no significantly enriched GO pathways were identified among the downregulated transcripts under hypoxia exposure (Supplementary Figure 3A). However, three GO pathways related to the extracellular region, extracellular space and enzyme regulator activity were significantly upregulated in the digestive gland of hypoxia-exposed *L. capensis* compared to the normoxic control (p_adj_<0.05) (Supplementary Figure 3B). No significantly upregulated or downregulated KEGG pathways were detected in the digestive gland transcriptome under hypoxia relative to control conditions (p_adj_>0.05) (Supplementary Figure 3C, D). After 24 h of post-hypoxia recovery, no significantly enriched down- or upregulated GO or KEGG pathways were found in the digestive gland compared to normoxic controls (p_adj_>0.05) (Supplementary Figure 4). The lysosome pathway was the only one found to be enriched in the digestive gland after 24 h of recovery compared to the hypoxia state, and it was downregulated (p_adj_<0.05) (Supplementary Figure 5).

### Reactome metabolic pathway enrichment analysis

Reactome analysis conducted on eukaryotic DEGs found strong evidence (FDR<0.05) for downregulation of 42 metabolic pathways in the gills of hypoxia-exposed *L. capensis* (Supplementary Table 5). It should be noted that because human homologs were used for Reactome pathway inferences, some identified reactions and pathways do not have direct one-to-one relationships between human and bivalve physiological processes, and instead, should be considered as proxies for underlying essential cellular and molecular interactions. Specifically, we found that the downregulated pathways in hypoxia were related to metabolism of proteins and RNA, cellular responses to stress (amino acid deficiency), metabolism, nervous system development, and infectious (particularly viral) disease. In such multi-species context, the individual genes involved in viral-related pathways (such as SARS-CoV-related ones) can be interpreted as putative players in a broad anti-viral or anti-microbial bivalve immune response and/or response against double-stranded RNAs (Rey-Campos et al., 2024) rather than precise molluscan response to SARS-CoV-1/-2 infections. Eighteen of these metabolic pathways showed strong overrepresentation of the downregulated DEGs, with >50% of the total genes in the respective pathway downregulated in hypoxia in the gills of *L. capensis*. These included multiple pathways associated with nonsense mediated decay (NMD) of mRNA, ribosome biogenesis and regulation, protein translation, viral-host interactions, and selenoamino acid metabolism (Supplementary Table 5). Other significantly downregulated pathways in hypoxia (FDR<0.05) showed less than 50% of the total genes in a pathway downregulated and were likewise related to ribosome biogenesis and regulation, protein translation, and viral-host interactions (Supplementary Table 5). Furthermore, Reactome pathways related to cellular stress response, starvation, metabolism, and neural development were significantly downregulated (FDR<0.05) (Supplementary Table 5).

Additional six pathways showed moderate evidence (0.05<FDR<0.1) for downregulation in the gills of the hypoxia-exposed *L. capensis*. Interestingly, three of these pathways were related to mitochondrial OXPHOS, including R-HAS-163200 (respiratory electron transport, ATP synthesis by chemiosmotic coupling, and heat production by uncoupling proteins), and R-HSA-6799198 (Complex I biogenesis) Reactome pathways, with 9-17 downregulated DEGs (corresponding to 12-16% of all genes in the respective pathway) (Supplementary Table 5). Reactome analysis identified a single pathway that showed moderate evidence of upregulation in the gills of hypoxia-exposed *L. capensis* (the citric acid (TCA) cycle and respiratory electron transport, FDR=0.06) (Supplementary Table 5). Notably, there was no evidence for alteration of glycolysis (Reactome pathway R-HSA-70171) in the gills or the digestive gland of the clams.

Metabolic pathways in the digestive gland of *L. capensis* were less affected by hypoxia exposure than those in the gills (Supplementary Table 6). No pathways were significantly enriched for downregulated genes in the digestive gland of hypoxia-exposed clams (FDR>0.1), and only three pathways related to membrane receptor-mediated signaling and immune function showed moderate evidence of upregulation (0.05<FDR<0.1) (Supplementary Table 6).

Despite the large number of DEGs found in the gills of *L. capensis* during post-hypoxia recovery relative to both hypoxia and the normoxic control, no overrepresentation of these genes in specific metabolic pathways was identified by Reactome (FDR>0.1). Similarly, the DEGs up- and downregulated in the digestive gland after 24 h of recovery relative to the control were not significantly overrepresented in any of the Reactome pathways (FDR>0.1). However, in the digestive gland the downregulated DEGs after 24 h of reoxygenation relative to hypoxia (FDR<0.1) were overrepresented in nine pathways associated with extracellular matrix maintenance and immune function (Supplementary Table 6). No pathway overrepresentation of the upregulated DEGs after 24 h of recovery relative to hypoxia was identified by the Reactome in the digestive gland (FDR>0.1).

### Metabolite profiles

Of the 35 metabolites quantified in the gills and digestive gland of *L. capensis* (see Zenodo DOI 10.5281/zenodo.15268761 for the full list of metabolites), six showed significant effects of the oxygen regime in the gills (p<0.05). Three amino acids (Met, Arg and Leu) accumulated during hypoxia exposure in the gill and gradually decreased to the baseline levels during reoxygenation (Fig. 7A-C). Similarly, AMP accumulated in *L. capensis* gills during hypoxia and returned back to the normoxic baseline after 1 h of reoxygenation (Fig. 7D). Succinate accumulated in the gills of *L. capensis* during hypoxia and continued to increase during the 1^st^ hour of reoxygenation, declining only after 24 h of post-hypoxia recovery (Fig. 7E). Malate concentration in the gills moderately increased during hypoxia and continued to rise throughout the 1–24 hour post-hypoxia recovery period (Fig. 7F).

**Figure 7.**
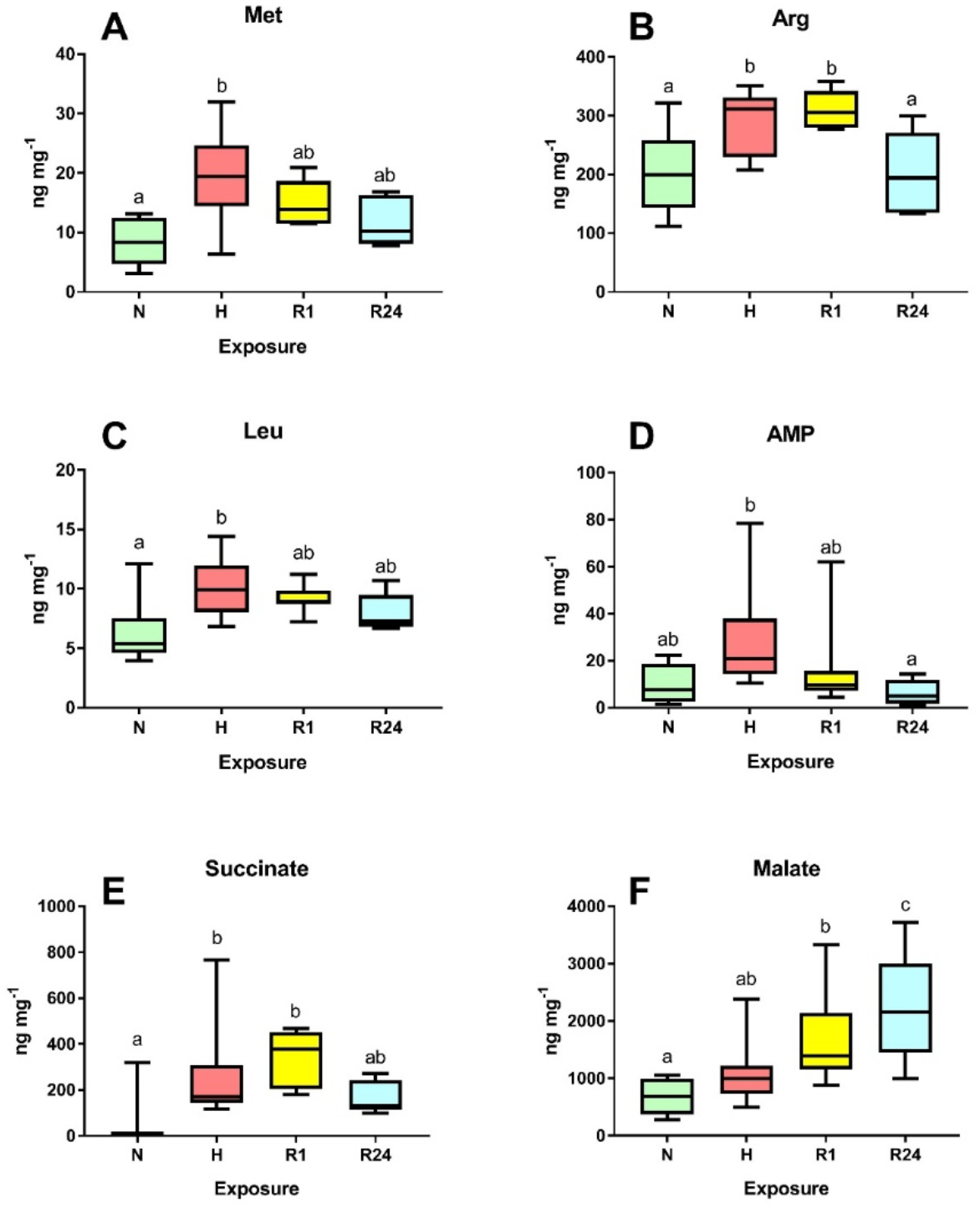
Effect of different oxygen conditions on concentrations of selected metabolites in the gills of *L. capensis*. Only metabolites that show significant effects of oxygen conditions (p<0.05) are shown. Different letters indicate significant differences between the values for 600 the respective exposure groups 601 (p<0.05). Conditions: N – normoxia, 602 H – hypoxia, R1 – 1 h reoxygenation, 603 R24 – 24 h reoxygenation. N=5-11 604 per group.

In the digestive gland, only S-adenosylmethionine (SAM) content showed significant differences between exposure groups (p<0.05). SAM concentrations in the digestive gland decreased in hypoxia and were partially restored after 24 h or reoxygenation (Fig. 8).

**Figure 8.**
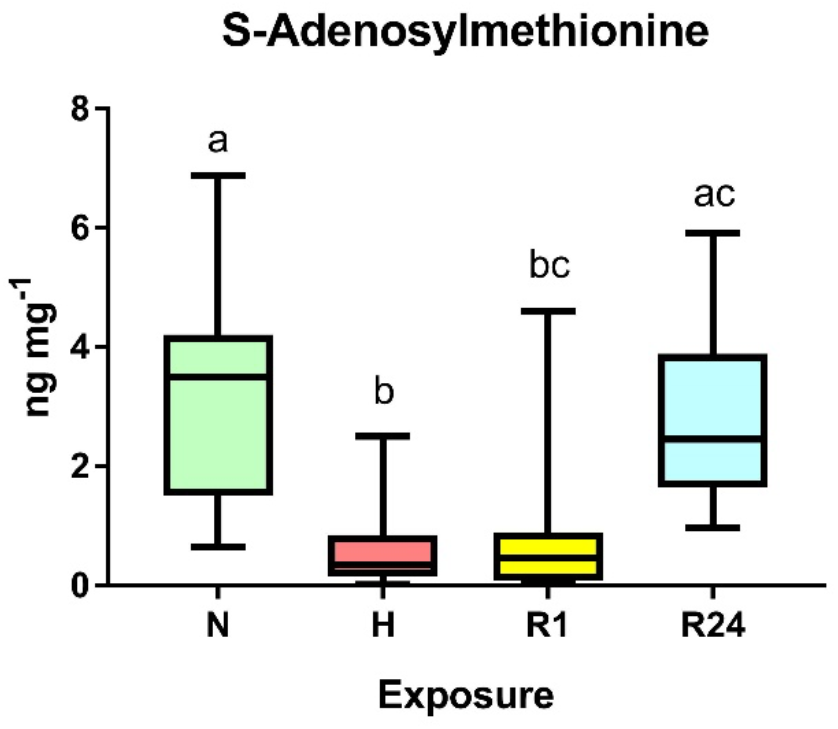
Effect of different oxygen conditions 613 on concentrations of SAM in the digestive 614 gland of *L. capensis*. Different letters indicate 615 significant differences between the values for 616 the respective exposure groups (p<0.05). 617 Conditions: N – normoxia, H – hypoxia, R1 – 618 1 h reoxygenation, R24 – 24 h reoxygenation. 619 N=7-11 per group.

The sPLS-DA analysis revealed a clear separation of metabolite profiles in the gills between clams from all exposure groups and the normoxic control (Fig. 9A). Metabolite profiles at early recovery (1h) closely resembled those of the hypoxia-exposed groups, but after 24 hours of recovery, they shifted towards the normoxic control profiles. In contrast, there was considerable overlap in metabolite profiles among the different oxygen exposure groups in the digestive gland, suggesting a less distinct metabolic response to oxygen variations in this tissue (Fig. 9B).

**Figure 9.**
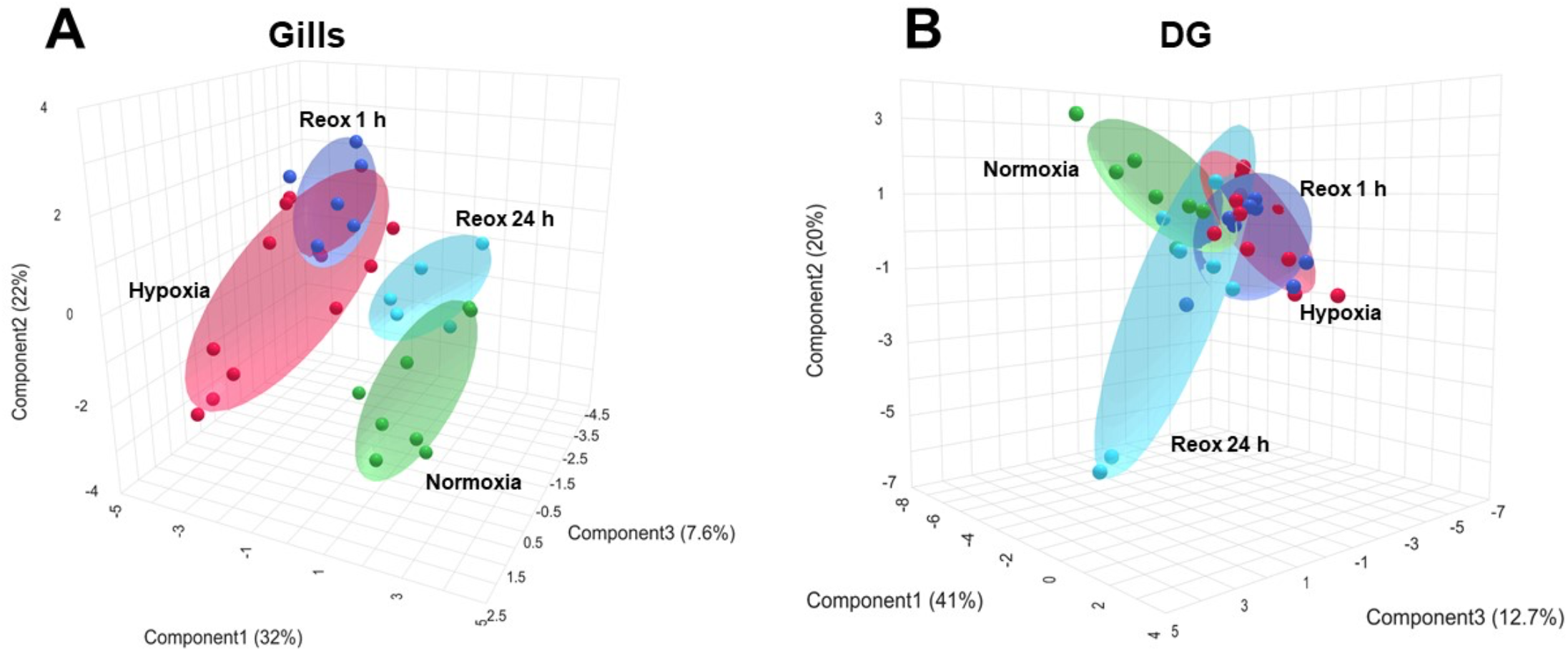
sPLS-DA 3D plots of the metabolite profiles of *L. capensis* gills (A) and the digestive gland (B) exposed to different oxygen conditions. Individual samples (shown by dots) are plotted in the space of three first principal components (PC). Numbers in brackets show the percent of the data variance explained by a certain PC. Shaded ellipsoids show 95% confidence intervals.

Metabolome-based pathway enrichment analysis did not reveal any significantly enriched metabolic pathways between the normoxic and hypoxic gills of *L. capensis*, or between the normoxic gills and those after 24 hours of reoxygenation (FDR > 0.1). However, significant alterations were observed in the TCA cycle (FDR < 0.05) as well as in pyruvate metabolism, butanoate metabolism, and cysteine and methionine metabolism (FDR<0.1) in the gills during the first hour of post-hypoxia recovery (Supplementary Table 7). In the digestive gland, no significantly enriched metabolic pathways were detected in either hypoxia or post-hypoxia recovery compared to the normoxic control (FDR > 0.1).

## Discussion

### Hypoxia response in *L. capensis* gill involves metabolic and immune suppression

The OMZ bivalve *L. capensis* exhibited a significant transcriptional response to hypoxia, with marked differences between the gills and the digestive gland. The gill transcriptome was considerably more responsive, showing 9,003 differentially expressed genes (DEGs) between normoxic and hypoxic conditions, compared to 2,697 DEGs in the digestive gland. However, only a small portion (~14–20%) of these DEGs could be assigned a putative function based on sequence homology with available gene and protein libraries, likely reflecting the high divergence of this deep-sea lucinid genome (Shin et al., 2024; Taylor et al., 2016; Taylor et al., 2011).

When focusing on genes with identified functions, the hypoxia-induced transcriptomic shift remained more pronounced in the gills (1,234 DEGs) than in the digestive gland (549 DEGs). While this difference may be partly influenced by the smaller sample size available for the transcriptomic analysis of the digestive gland, the metabolomic data also showed a lower level of hypoxia-induced change in this tissue, supporting the interpretation that the digestive gland is less sensitive to oxygen fluctuations than the gills. The gills serve as the primary organ for gas exchange and feeding in bivalves (Carroll and Catapane, 2007; Morton, 1960) and their function is highly sensitive to oxygen availability. Under severe hypoxia, shell closure stops oxygen and food intake, and as the limited oxygen within the shell is rapidly depleted, the bivalves transition to anaerobic metabolism (Brinkhoff et al., 1983; Ortmann and Grieshaber, 2003).

Anaerobic metabolism poses several challenges to animals, including energy deficiency due to the significantly lower ATP yield of anaerobic glycolysis compared to aerobic pathways, accumulation of potentially toxic metabolic byproducts, and metabolic acidosis (Brinkhoff et al., 1983; Hochachka and Mommsen, 1983; Hochachka and Mustafa, 1972; Pörtner et al., 1984). A key survival strategy in anoxia-tolerant animals is the suppression of metabolic rate, which involves a coordinated reduction in both ATP production and consumption (Hochachka et al., 1996; Hochachka and Guppy, 1987). Protein turnover is one of the most energy-demanding cellular processes (Li et al., 2014; Sokolova, 2021), and our study demonstrates that the downregulation of protein synthesis during hypoxia is a crucial energy-saving mechanism in the gills of *L. capensis*. GO, KEGG enrichment, and Reactome analyses revealed a coordinated downregulation of protein transcription, translation, and transport, along with suppressed metabolism of amino acids and their derivatives—critical for maintaining a balanced amino acid supply for protein synthesis—in the hypoxic gills of *L. capensis*. Interestingly, the pathway typically activated in response to amino acid deprivation—such as the cellular response to starvation (e.g., Reactome R-HSA-9711097)—was also downregulated, likely reflecting an adaptive energy-conservation strategy that prioritizes essential survival functions like cellular homeostasis over energetically costly responses to amino acid depletion. Similarly, energy-demanding processes involved in neural development (e.g., SLIT and ROBO signaling, axon guidance) were downregulated, possibly reflecting either an energy-saving mechanism or a secondary effect of the broader suppression of protein synthesis.

Although pathway enrichment analyses did not identify proteolysis or protein catabolic processes among the significantly enriched GO terms in *L. capensis* gills, a substantial proportion of DEGs—62 downregulated and 25 upregulated, comprising 12% and 11% of the total downregulated and upregulated DEG pools, respectively—were associated with proteolysis. Although the transcriptomic response of the digestive gland to hypoxia was weaker, with fewer DEGs and significantly enriched pathways compared to the gill, the regulation of protein synthesis (18 downregulated DEGs, representing 11% of the altered transcriptome) and proteolysis (11 upregulated and 38 downregulated DEGs, accounting for 6% and 10% of the altered transcriptome, respectively) was prominent. Notably, proteolysis-related genes constituted the largest or second-largest functional category of DEGs in both the gill and digestive gland transcriptomes. The apparent discrepancy between the abundance of proteolysis-related DEGs and the lack of significant GO term enrichment likely results from the wide distribution and heterogeneity of these genes across various sub-processes, thereby diluting their enrichment signal. Nevertheless, the prominence of proteolysis-related DEGs is biologically significant in the context of hypoxia, where a shutdown of protein synthesis would require the extended functionality of existing proteins (Hand and Hardewig, 1996). Simultaneously, the efficient removal of proteins damaged by hypoxic stress is essential for maintaining cellular integrity and function (Agrawal et al., 2017; Fawcett et al., 2015). Consequently, the regulation of proteolytic pathways is essential for balancing protein turnover during metabolic suppression in hypoxic tissues of *L. capensis*, even if this is not captured in pathway-level enrichment analyses.

Intriguingly, a substantial proportion of downregulated genes in the gills of *L. capensis* were associated with immune functions (62 DEGs, 8%). In contrast, this immune suppression was not evident in the digestive gland, where only 7 immune-related genes (4%) were downregulated, while 62 (16%) were upregulated. Reactome analysis further indicated enrichment of these downregulated immune genes in pathways linked to viral-host interactions in the gills. This gill-specific transcriptional suppression of immune pathways likely represents an energy-conserving adaptation to hypoxic stress, rather than a response to an active immune challenge. Notably, similar patterns of immune gene modulation under hypoxia have been observed in other bivalves, including the shallow-water mytilids *Mytilus edulis* and *M. chilensis*, as well as the arcid clam *Tegillarca granosa* (Cheng et al., 2024; Montúfar-Romero et al., 2024; Wu et al., 2024a), suggesting that hypoxia-driven immunomodulation may be a conserved response across Bivalvia.

The enrichment of downregulated transposition-related pathways and transposable element dynamics (58 DEGs, 7%) in the gills of *L. capensis* suggests a broader strategy to maintain genomic integrity in response to environmental fluctuations. Transposable elements (TEs) make up a significant portion of all genomes, accounting for anywhere between 3% and 50% of the genome’s content, varying by species (Capy et al., 2000). Although the functions of TEs are not yet fully understood, studies in plants and animals indicate that environmental stressors like temperature, UV radiation, oxidative stress or nutritional stress can mobilize TEs (Capy et al., 2000; de Oliveira et al., 2021; Pappalardo et al., 2021). It has been proposed that stress-activated TEs play an adaptive role by increasing mutation rates and genetic variability, spreading regulatory elements to form stress-inducible gene networks and providing material for natural selection to develop traits that help species cope with stress (Capy et al., 2000; Pappalardo et al., 2021). However, the regulation of TE dynamics in response to stress is not uniform; different TE families can be either activated or repressed, with the degree of activation or suppression varying according to the nature and intensity of the stressor (de Oliveira et al., 2021).

Research on the effects of hypoxia on TE dynamics remains limited. Nevertheless, studies in mammalian cells have shown that hypoxia suppresses TE transcription (Park et al., 2023). Additionally, artificial derepression of TEs under hypoxic conditions has been found to hyperactivate immune-inflammatory responses, induce DNA damage, and ultimately lead to cell death (Park et al., 2023). Thus, the suppression of TE dynamics in the gills of *L. capensis* may play a crucial role in maintaining genomic stability, preventing the activation of immune responses, and supporting cell viability under hypoxic conditions. The coordinated downregulation of TE-related and immune transcripts, particularly in the gills, may also mitigate the inadvertent loss of essential symbionts due to hyperactivation of immune mechanisms. Moreover, this downregulation may reflect an energy-conserving strategy in which metabolic processes are prioritized over immune and genomic defense mechanisms in response to oxygen limitation. In summary, the simultaneous downregulation of immune responses and mobile genetic element dynamics in the gills of *L. capensis* suggests an evolutionary strategy for optimizing energy allocation under hypoxic conditions while ensuring the maintenance of symbionts and genomic resilience in an unstable environment.

As expected, the host’s oxidative phosphorylation (OXPHOS) system was suppressed in the hypoxic gills of *L. capensis*. This suppression involved the downregulation of the respiratory electron transport, ATP synthesis via chemiosmotic coupling, uncoupling proteins (UCPs), and Complex I biogenesis (Reactome pathways R-HSA-163200, R-HSA-611105 and R-HSA-6799198). Given that Complex I is a significant source of reactive oxygen species (ROS) (Brondani et al., 2023; Chouchani et al., 2016; Sokolov et al., 2021), its downregulation may serve to mitigate oxidative stress, particularly during the reoxygenation phase. Notably, we found no evidence of transcriptional upregulation of antioxidant pathways in the gills or digestive gland of *L. capensis*, neither during hypoxia nor reoxygenation. Instead, selenoamino acid metabolism, including selenocysteine synthesis, was significantly downregulated in the hypoxic gills. Selenocysteine, often referred to as the 21^st^ amino acid, is incorporated into selenoproteins, such as glutathione peroxidases and thioredoxin reductases, which play key roles in antioxidant defense and redox regulation (Steinbrenner et al., 2016). The reduced transcription of selenoprotein-related pathways suggests diminished demand for antioxidant activity in hypoxia-exposed clams. This hypothesis is supported by earlier reports that reoxygenation in bivalve mitochondria does not cause a significant increase in ROS efflux and that these mitochondria can effectively oxidize succinate with minimal reverse electron transport (RET) through Complex I (Adzigbli et al., 2024; Adzigbli et al., 2022; Sokolov et al., 2021; Steffen et al., 2021; Steffen et al., 2023). Additionally, comparative studies across bivalve species with varying thermal stress tolerances have shown that basal levels and inducibility of antioxidant enzymes do not correlate with thermal tolerance (Dowd and Somero, 2023; Falfushynska et al., 2016). These findings suggest that *L. capensis* and other stress-tolerant bivalves rely more on regulating (or tolerating) ROS production, rather than upregulating antioxidant defenses, as a survival strategy. This adaptive approach may also serve to conserve energy by minimizing the need for de novo protein synthesis, including that of antioxidants, during hypoxia.

The gills of *L. capensis* harbor sulfur-oxidizing symbionts (*Candidatus Thiodiazotropha*) that detoxify H_2_S and support the host by fixing CO_2_ and assimilating ammonium under hypoxia (Amorim et al., 2022; Cary and Felbeck, 1989; Yuen et al., 2019). In this study, we observed upregulation of prokaryotic transcripts associated with sulfur metabolism, CO_2_ fixation, electron transport, and protein synthesis—potentially reflecting enhanced symbiont activity related to detoxification and energy production during hypoxia. However, interpretation of the bacterial signal is constrained by the use of poly-A enrichment, which biases against non-polyadenylated microbial RNA, and the taxonomic resolution of the data does not allow for definitive attribution of transcripts to *Ca. Thiodiazotropha*. Despite these limitations, the functional signatures detected in the gills are consistent with the established roles of chemosynthetic symbionts in lucinid clams and provide preliminary insight into their transcriptional responses to low oxygen conditions.

### Hypoxic digestive gland of *L. capensis* upregulates structural maintenance

In comparison to the gill, the transcriptomic response of the digestive gland to hypoxia was notably muted (Fig. 3, Supplementary Table 4). This attenuated response, relative to that of the gills, may be attributed to the digestive gland’s distinct physiological role. While the gills are directly involved in oxygen uptake, the digestive gland primarily functions in digestion and nutrient storage, activities not immediately related to gas exchange (Lobo-da-Cunha, 2019). Moreover, as an internal organ, the digestive gland may encounter hypoxic conditions in a delayed or buffered manner compared to the gills. Transcriptomic analyses in the Pacific oyster, *Crassostrea gigas*, revealed differential expression of prolyl hydroxylase domain-containing protein 2 (PHD2) isoforms between the gills and the digestive gland, with the gills expressing a more oxygen-sensitive, strongly inducible, and active isoform (Meng et al., 2022). These findings suggest that the reduced sensitivity of the digestive gland transcriptome to hypoxia reflects both the functional specialization and molecular composition of the oxygen-sensing pathways in this organ, which make it less responsive to fluctuations in oxygen availability.

In the digestive gland transcriptome of *L. capensis* exposed to 36 h of severe hypoxia, no significantly downregulated pathways were observed. However, two pathways related to extracellular matrix (ECM) formation and tissue maintenance, mediated by fibroblast growth factor receptor (FGFR) 2 activation, were upregulated. While the role of FGFR in mollusks is not well understood, in mammals, FGFR signaling regulates cell proliferation, differentiation, and tissue repair (Xie et al., 2020). The upregulation of FGFR signaling in *L. capensis* likely reflects an increased investment in tissue maintenance and repair under hypoxia. This hypothesis is supported by the analysis of manually annotated genes showing upregulation of genes involved in ECM formation and maintenance, representing 8% of the upregulated transcriptome (32 DEGs), including those regulating collagen metabolism, proteoglycan synthesis, as well as cellular adhesion (16 DEGs, 4% of the upregulated transcriptome). These findings suggest a focus on preserving the structural integrity of the digestive gland and its epithelium. Additionally, the observed upregulation of immune-related genes (62 DEGs, 16% of the upregulated transcriptome), particularly those involved in the trafficking and processing of endosomal Toll-like receptors (TLRs) (Reactome pathway R-HSA-1679131), further supports this interpretation. TLRs are critical in tissue repair and regeneration, as they respond to pathogen-associated molecular patterns (PAMPs) released by the microbiome or damage-associated molecular patterns (DAMPs) from dying cells, triggering tissue recovery processes (Ioannou and Voulgarelis, 2010). Studies on the mud crab *Scylla paramamosain* have demonstrated a similar role for FGFR signaling in regulating the TLR pathway (Li et al., 2022), suggesting that this function is conserved across invertebrates and vertebrates. These findings suggest that the hypoxic response in the digestive gland of *L. capensis* prioritizes tissue integrity through ECM and epithelial maintenance, supported by the absence of significant downregulation of protein synthesis in this organ.

### Metabolome stability during oxygen fluctuations in *L. capensis*

Despite substantial transcriptomic reorganization, the tissue metabolome of both the gill and digestive gland of *L. capensis* remained remarkably stable during oxygen fluctuations. Of the 35 quantified metabolites in the gill, only five showed significant changes under hypoxia, and three in response to reoxygenation, compared to the normoxic baseline. Succinate accumulation in hypoxic gills, a typical response in marine bivalves, indicates a shift to mitochondrial anaerobiosis (Pörtner et al., 1984; Tielens et al., 2002). Notably, no alanine accumulation was observed, suggesting that cytosolic anaerobiosis is either bypassed or transient in this hypoxia-adapted species. Elevated AMP levels reflect an imbalance between ATP breakdown and resynthesis, reducing cellular energy charge, a trend quickly reversed upon reoxygenation, along with a malate overshoot, indicating TCA reactivation. Three amino acids (Met, Leu, Arg) accumulated under hypoxia, returning to baseline after reoxygenation (Met, Leu at 1h; Arg at 24h). Arginine accumulation likely reflects phosphagen breakdown for anaerobic ATP production (Livingstone, 1991), while Met and Leu accumulation might be due to a hypoxia-induced shutdown of protein synthesis.

In the digestive gland, the only metabolite exhibiting a significant change under hypoxia was a substantial (~5.4-fold) decrease in S-adenosylmethionine (SAM) content. This reduction suggests a diminished methylation capacity, potentially impacting DNA and protein methylation (Loenen, 2006). Given SAM’s involvement in numerous cellular functions, it is challenging to identify which specific processes in the digestive gland of *L. capensis* are affected by this decline. However, the rapid restoration of SAM levels upon reoxygenation, returning to normoxic levels within 24 hours, suggests that any associated effects are likely transient. Notably, there was no evidence of upregulation in anaerobic ATP production pathways in the digestive gland, as indicated by the lack of succinate or alanine accumulation. This is consistent with transcriptomic findings, which highlight the digestive gland’s low sensitivity to oxygen fluctuations, and suggests that its ATP demand during 36 h of hypoxia is met either by residual aerobic metabolism or by hemolymph-mediated transport of energy and nutrients from symbiont-containing tissues, such as the gill (Frizzo et al., 2021).

When comparing the hypoxia-induced metabolome shifts of *L. capensis* with other marine bivalves, including hypoxia-tolerant intertidal species, *L. capensis* demonstrates remarkable metabolic resilience. In *M. edulis*, for instance, 24 h of severe hypoxia caused significant shifts in 19 out of 20 amino acids in both the gills and whole body, with most amino acids accumulating, while others, such as aspartate (utilized in anaerobic alanine production), were depleted (Haider et al., 2020; Wu et al., 2024b). Similarly, in the Pacific oyster *Crassostrea gigas*, hypoxia led to the accumulation or depletion of several amino acids - 7 out of 20 after 24 h and 15 out of 20 after 6 days (Haider et al., 2020). Furthermore, *M. edulis* experienced hypoxia-induced changes in metabolic pathways related to energy, carbohydrate, amino acid, and co-factor metabolism (Wu et al., 2024b). In *C. gigas*, intermittent hypoxia affected multiple key pathways, including alanine, aspartate, and glutamate metabolism, arginine biosynthesis, metabolism of aromatic amino acids, and the TCA cycle (Bruhns et al., 2023). Even mild hypoxia (2 mg L^−1^ O_2_) in the pearl oyster *Pinctada fucata martensii* triggered a substantial reorganization of 25 metabolic pathways, including metabolism of tRNA, amino acids, glycerophospholipids, and ABC transporters activity (Yang et al., 2023). In contrast, the metabolome of *L. capensis* in both the gill and digestive gland remained notably stable during hypoxia and reoxygenation, underscoring its exceptional resilience to oxygen fluctuations.

### Recovery: A return to baseline or a distinct metabolic state?

Pathway enrichment analyses revealed that immunosuppression initiated during hypoxic exposure in the gills of *L. capensis* intensified during the first hour of reoxygenation, as evidenced by the suppression of multiple immune-related KEGG pathways associated with immune receptor signaling, chemokine signaling, and PAMP/DAMP sensing. Additionally, inflammation signaling pathways were significantly suppressed during the first hour of reoxygenation compared to the hypoxic state. After 24 h of reoxygenation, no immunity-related pathways were significantly enriched relative to the normoxic baseline or the hypoxic state, despite the identification of 29 downregulated and 11 upregulated immune-related DEGs compared to the baseline, indicating gradual return to the normoxic baseline state.

Differential regulation of gill proteolysis during the recovery phase indicates a significant shift in cellular priorities in response to reoxygenation. In the first 1 h of recovery, there were 49 downregulated DEGs related to proteolysis, representing 12% of the total downregulated pool, alongside 27 upregulated DEGs, accounting for 9% of the upregulated pool relative to the normoxic baseline. This suggests that the gill is prioritizing the degradation of specific proteins during this initial recovery phase. This trend continues after 24 h of reoxygenation, with 23 upregulated and 27 downregulated DEGs related to proteolysis, representing 9% and 8% of their respective pools. Despite the significant representation of proteolysis-related DEGs in the recovering gill transcriptome, the absence of specific enriched pathways indicates that these genes are widely distributed among various pathways and subprocesses. In contrast, only 23 and 20 downregulated DEGs related to protein synthesis were identified after 1 and 24 hours of recovery, respectively, with 12 upregulated DEGs present at both recovery phases. This contrasts sharply with the hypoxic state, during which 150 downregulated and 12 upregulated DEGs were observed. Moreover, two GO pathways - chromosome binding and histone binding - were significantly enriched among the upregulated genes in *L. capensis* gills after 24 hours of recovery, suggesting an enhancement in transcriptional regulation. Collectively, these findings indicate a rapid restoration of proteosynthetic activity as the gills transition from a hypoxic to a reoxygenated state.

Overall, reoxygenation signifies a return of many essential cellular processes to the normoxic baseline, with a remarkably rapid restoration of protein synthesis and a more gradual recovery of other processes, such as protein degradation and immune function. The number of downregulated host’s DEGs decreases progressively from hypoxia (819 DEGs) to 1 h of recovery (409 DEGs) and then to 24 h of recovery (261 DEGs). This trend indicates a depression of processes that were halted during hypoxia as part of a metabolic suppression strategy. Despite the gradual decrease in downregulated DEGs, the gill transcriptome reveals a significant upregulation of various processes during recovery, with 289 and 328 DEGs exceeding the normoxic baseline after 1 and 24 hours of reoxygenation, respectively. This upregulation is not merely a carryover from the hypoxic state; instead, distinct functional categories of DEGs are often overrepresented in recovery compared to hypoxia. The overshoot in TCA cycle activity shown by metabolomic data further suggests that recovery involves the active regulation of specific metabolic processes rather than a simple return to baseline levels. In hypoxia-tolerant organisms, recovery is frequently associated with enhanced metabolism to restore homeostasis, manifested at the organismal level as an increase in oxygen consumption, commonly referred to as “oxygen debt” (Lewis et al., 2007; Van den Thillart and Verbeek, 1991; Vismann and Hagerman, 1996). Our study shows that the molecular mechanisms contributing to this spike in metabolism involve stimulation of multiple cellular processes that span broad metabolic networks and thus do not prominently feature in specific enriched pathways during the recovery of gills relative to the normoxic steady state.

Recovery in the digestive gland was less pronounced, consistent with the lower transcriptomic impact of hypoxia in this tissue compared to the gills. Both hypoxia and 24 hours of recovery showed similar numbers of DEGs, with 167 downregulated and 382 upregulated in hypoxia, and 144 downregulated and 384 upregulated after 24 h of recovery. The only significantly enriched pathway for downregulated DEGs in the recovering digestive gland was the lysosome pathway, suggesting a suppression of intracellular digestion - likely as an energy-saving strategy to minimize specific dynamic action (Goodrich et al., 2024). Despite these shifts, the digestive gland demonstrated considerable transcriptomic and metabolic stability during oxygen fluctuations, reflecting its resilience and lower sensitivity to hypoxia compared to other tissues like the gills.

## Conclusions

This study provides the first comprehensive insights into the metabolic regulation of the OMZ-adapted bivalve *L. capensis* in response to fluctuating oxygen conditions. The data reveal pronounced organ-specific specialization, with the gills exhibiting strong transcriptional responses to oxygen variability, in stark contrast to the relative transcriptional stability observed in the digestive gland.

Under hypoxic conditions, the *L. capensis* gills exhibited a coordinated downregulation of key metabolic processes, including protein synthesis, transposable element activity, and immune function. This suggests a tightly regulated energy conservation strategy and potential mechanisms to maintain symbiont stability and genomic integrity. Additionally, there was evidence for the activation of endosymbiont metabolism, consistent with its well-established role in host energy acquisition and sulfide detoxification during hypoxia (Amorim et al., 2022; Cary and Felbeck, 1989; Yuen et al., 2019). In contrast, the digestive gland exhibited minimal transcriptional shifts during anoxia. Upregulation of specific pathways suggests an investment in structural maintenance, which likely contributes to the observed stability of the metabolome under low oxygen conditions.

Reoxygenation induced an active and asymmetric recovery trajectory in the gill, characterized by a rapid restoration of protein synthesis and a more gradual normalization of protein degradation and immune-related processes. The overshoot observed in TCA cycle intermediates, alongside the derepression of pathways previously downregulated during hypoxia, indicates that reoxygenation involves active metabolic reprogramming rather than a simple return to baseline.

Together, these findings highlight the organ-level metabolic specialization that underpin the resilience of *L. capensis* to OMZ dynamics. This work provides essential groundwork for understanding the molecular strategies enabling benthic invertebrates to persist in increasingly oxygen-depleted marine environments.

## Acknowledgements

This study was part of the EVAR project funded by the German Federal Ministry of Education and Research (grant no.: 03V01279) awarded to MZ. We gratefully acknowledge the support of the Mare Balticum Fellowship Program of the University of Rostock, which enabled HP to establish research collaboration related to this work. The LC-MS/MS equipment used in this study was financed through a grant from the Hochschul-Bau-Förderungsgesetz (HBFG) program (award no.: GZ INST 264/125-1FUGG). We thank Christian Burmeister for his assistance with the seawater analyses. Open Access funding was provided by the University of Rostock. The transcriptomic data have been submitted to the NCBI GenBank under the BioProject accession number PRJNA1244758. All other data that support the findings of this study are available in the Supplementary Materials and as Zenodo datasets at https://zenodo.org/, DOI 10.5281/zenodo.15268761 and 10.5281/zenodo.16367847.

## Data Availability

The transcriptomic data have been deposited in NCBI GenBank Sequence Read Archive (SRA) under BioProject accession number PRJNA1244758. The metabolomic dataset and the dataset related to the transcriptomic analyses are publicly available on Zenodo at 10.5281/zenodo.15268761 and 10.5281/zenodo.16367847, respectively, under the Creative Commons Attribution Share Alike 4.0 International License.

## References

Adzigbli L, Ponsuksili S, Sokolova I. Mitochondrial responses to constant and cyclic hypoxia depend on the oxidized fuel in a hypoxia-tolerant marine bivalve Crassostrea gigas. Scientific Reports 2024; 14: 9658.

Adzigbli L, Sokolov EP, Ponsuksili S, Sokolova IM. Tissue- and substrate-dependent mitochondrial responses to acute hypoxia–reoxygenation stress in a marine bivalve (Crassostrea gigas). Journal of Experimental Biology 2022; 225.

Agrawal A, Rathor R, Suryakumar G. Oxidative protein modification alters proteostasis under acute hypobaric hypoxia in skeletal muscles: a comprehensive in vivo study. Cell Stress Chaperones 2017; 22: 429–443.

Altschul SF, Madden TL, Schäffer AA, Zhang J, Zhang Z, Miller W, et al. Gapped BLAST and PSI-BLAST: a new generation of protein database search programs. Nucleic Acids Research 1997; 25: 3389–3402.

Amorim K, Loick-Wilde N, Yuen B, Osvatic JT, Wäge-Recchioni J, Hausmann B, et al. Chemoautotrophy, symbiosis and sedimented diatoms support high biomass of benthic molluscs in the Namibian shelf. Sci Rep 2022; 12: 9731.

Amorim K, Piontkivska H, Zettler ML, Sokolov E, Hinzke T, Nair AM, et al. Transcriptional response of key metabolic and stress response genes of a nuculanid bivalve, Lembulus bicuspidatus from an oxygen minimum zone exposed to hypoxia-reoxygenation. Comparative Biochemistry and Physiology Part B: Biochemistry and Molecular Biology 2021; 256: 110617.

Amorim K, Zettler ML. Gradients and instability: Macrozoobenthic communities in the Benguela Upwelling System off Namibia. Estuarine, Coastal and Shelf Science 2023; 291: 108421.

Benjamini Y, Hochberg Y. Controlling the False Discovery Rate: A Practical and Powerful Approach to Multiple Testing. Journal of the Royal Statistical Society: Series B (Methodological) 1995; 57: 289–300.

Breitburg D, Baumann H, Sokolova I, Frieder C. 6. Multiple stressors-forces that combine to worsen deoxygenation and its effects. Ocean deoxygenation: Everyone’s problem. Causes, impacts, consequences and solutions. IUCN, Gland, Switzerland, 2019, pp. 225–247.

Brinkhoff W, Stöckmann K, Grieshaber M. Natural occurrence of anaerobiosis in molluscs from intertidal habitats. Oecologia 1983; 57: 151–155.

Brondani M, Roginski AC, Ribeiro RT, de Medeiros MP, Hoffmann CIH, Wajner M, et al. Mitochondrial dysfunction, oxidative stress, ER stress and mitochondria-ER crosstalk alterations in a chemical rat model of Huntington’s disease: Potential benefits of bezafibrate. Toxicol Lett 2023; 381: 48–59.

Bruhns T, Timm S, Feußner N, Engelhaupt S, Labrenz M, Wegner M, et al. Combined effects of temperature and emersion-immersion cycles on metabolism and bioenergetics of the Pacific oyster Crassostrea (Magallana) gigas. Mar Environ Res 2023; 192: 106231.

Capy P, Gasperi G, Biémont C, Bazin C. Stress and transposable elements: co-evolution or useful parasites? Heredity 2000; 85: 101–106.

Carroll MA, Catapane EJ. The nervous system control of lateral ciliary activity of the gill of the bivalve mollusc, Crassostrea virginica. Comp Biochem Physiol A Mol Integr Physiol 2007; 148: 445–50.

Cary S, Felbeck H. Habitat characterization and nutritional strategies of the endosymbiont-bearing bivalve Lucinoma aequizonata. Marine Ecology-progress Series - MAR ECOL-PROGR SER 1989; 55: 31–45.

Chen J, Hu Z, Li P, Wang G, Wei H, Li Q, et al. Transcriptomic atlas for hypoxia and following re-oxygenation in Ancherythroculter nigrocauda heart and brain tissues: insights into gene expression, alternative splicing, and signaling pathways. Frontiers in Genetics 2024; 15.

Cheng H, Peng Z, Zhao C, Jin H, Bao Y, Liu M. The transcriptomic and biochemical responses of blood clams (Tegillarca granosa) to prolonged intermittent hypoxia. Comparative Biochemistry and Physiology Part B: Biochemistry and Molecular Biology 2024; 270: 110923.

Chouchani Edward T, Pell Victoria R, James Andrew M, Work Lorraine M, Saeb-Parsy K, Frezza C, et al. A Unifying Mechanism for Mitochondrial Superoxide Production during Ischemia-Reperfusion Injury. Cell Metabolism 2016; 23: 254–263.

Connor KM, Gracey AY. High-resolution analysis of metabolic cycles in the intertidal mussel Mytilus californianus. Am J Physiol Regul Integr Comp Physiol 2012; 302: R103–11.

Davidson NM, Oshlack A. Corset: enabling differential gene expression analysis for de novoassembled transcriptomes. Genome Biology 2014; 15: 410.

de Oliveira DS, Rosa MT, Vieira C, Loreto ELS. Oxidative and radiation stress induces transposable element transcription in Drosophila melanogaster. Journal of Evolutionary Biology 2021; 34: 628–638.

Deutsch C, Penn JL, Lucey N. Climate, Oxygen, and the Future of Marine Biodiversity. Annual Review of Marine Science 2024; 16: 217–245.

Dowd WW, Somero GN. Oxidative stress effects are not correlated with differences in heat tolerance among congeners of Mytilus. J Exp Biol 2023; 226.

Eisenbarth S, Zettler ML. Diversity of the benthic macrofauna off northern Namibia from the shelf to the deep sea. Journal of Marine Systems 2016; 155: 1–10.

Falfushynska HI, Phan T, Sokolova IM. Long-Term Acclimation to Different Thermal Regimes Affects Molecular Responses to Heat Stress in a Freshwater Clam Corbicula Fluminea. Scientific Reports 2016; 6: 39476.

Fawcett EM, Hoyt JM, Johnson JK, Miller DL. Hypoxia disrupts proteostasis in Caenorhabditis elegans. Aging Cell 2015; 14: 92–101.

Friedrich J, Janssen F, Aleynik D, Bange HW, Boltacheva N, Çagatay MN, et al. Investigating hypoxia in aquatic environments: diverse approaches to addressing a complex phenomenon. Biogeosciences 2014; 11: 1215–1259.

Frizzo R, Bortoletto E, Riello T, Leanza L, Schievano E, Venier P, et al. NMR Metabolite Profiles of the Bivalve Mollusc Mytilus galloprovincialis Before and After Immune Stimulation With Vibrio splendidus. Frontiers in Molecular Biosciences 2021; Volume 8 - 2021.

Goodrich HR, Wood CM, Wilson RW, Clark TD, Last KB, Wang T. Specific dynamic action: the energy cost of digestion or growth? Journal of Experimental Biology 2024; 227.

Grabherr MG, Haas BJ, Yassour M, Levin JZ, Thompson DA, Amit I, et al. Full-length transcriptome assembly from RNA-Seq data without a reference genome. Nat Biotechnol 2011; 29: 644–52.

Gracey AY, Chaney ML, Boomhower JP, Tyburczy WR, Connor K, Somero GN. Rhythms of gene expression in a fluctuating intertidal environment. Curr Biol 2008; 18: 1501–7.

Grasshoff K, Kremling K, Ehrhardt Me. Methods of Seawater Analysis. 3rd edition. Weinheim: John Wiley & Sons, 2009.

Haider F, Falfushynska HI, Timm S, Sokolova IM. Effects of hypoxia and reoxygenation on intermediary metabolite homeostasis of marine bivalves Mytilus edulis and Crassostrea gigas. Comparative Biochemistry and Physiology Part A: Molecular & Integrative Physiology 2020; 242: 110657.

Halliwell B, Gutteridge JMC. Free Radicals in Biology and Medicine. Oxford, New York: Oxford University Press, 1999.

Hand SC, Hardewig I. Downregulation of Cellular Metabolism During Environmental Stress: Mechanisms and Implications. Annual Review of Physiology 1996; 58: 539–563.

Hermes-Lima M. Oxygen in biology and biochemistry: role of free radicals. Functional metabolism: Regulation and adaptation. 1. Wiley-Liss Hoboken, 2004, pp. 319–366.

Hochachka PW, Buck LT, Doll CJ, Land SC. Unifying theory of hypoxia tolerance: molecular/metabolic defense and rescue mechanisms for surviving oxygen lack. Proc Natl Acad Sci U S A 1996; 93: 9493–8.

Hochachka PW, Guppy M. Metabolic arrest and the control of biological time. Cambridge (Ma), London (England): Harvard University Press, 227 pp., 1987.

Hochachka PW, Mommsen TP. Protons and anaerobiosis. Science 1983; 219: 1391–1397.

Hochachka PW, Mustafa T. Invertebrate Facultative Anaerobiosis. Science 1972; 178: 1056–1060.

Howarth R, Chan F, Conley DJ, Garnier J, Doney SC, Marino R, et al. Coupled biogeochemical cycles: eutrophication and hypoxia in temperate estuaries and coastal marine ecosystems. Frontiers in Ecology and the Environment 2011; 9: 18–26.

Ioannou S, Voulgarelis M. Toll-Like Receptors, Tissue Injury, and Tumourigenesis. Mediators of Inflammation 2010; 2010: 581837.

Kern B, Ivanina AV, Piontkivska H, Sokolov EP, Sokolova IM. Molecular characterization and expression of a novel homolog of uncoupling protein 5 (UCP5) from the eastern oyster Crassostrea virginica (Bivalvia: Ostreidae). Comp Biochem Physiol Part D Genomics Proteomics 2009; 4: 121–7.

Lane N. Oxygen: The Molecule That Made the World, 2002.

Lane N. Power, Sex, Suicide: Mitochondria and the Meaning of Life, 2005.

Lê Cao K-A, Boitard S, Besse P. Sparse PLS discriminant analysis: biologically relevant feature selection and graphical displays for multiclass problems. BMC Bioinformatics 2011; 12: 253.

Lewis JM, Costa I, Val AL, Almeida-Val VM, Gamperl AK, Driedzic WR. Responses to hypoxia and recovery: repayment of oxygen debt is not associated with compensatory protein synthesis in the Amazonian cichlid, Astronotus ocellatus. J Exp Biol 2007; 210: 1935–43.

Li B-W, Xu W-B, Dong W-R, Zhang Y-M, Cheng Y-X, Chen D-Y, et al. Identification and function analysis of two fibroblast growth factor receptor (FGFR) from Scylla paramamosain: The evidence of FGFR involved in innate immunity in crustacean. Fish & Shellfish Immunology 2022; 131: 602–611.

Li B, Dewey CN. RSEM: accurate transcript quantification from RNA-Seq data with or without a reference genome. BMC Bioinformatics 2011; 12: 323.

Li GW, Burkhardt D, Gross C, Weissman JS. Quantifying absolute protein synthesis rates reveals principles underlying allocation of cellular resources. Cell 2014; 157: 624–35.

Lim SJ, Davis BG, Gill DE, Walton J, Nachman E, Engel AS, et al. Taxonomic and functional heterogeneity of the gill microbiome in a symbiotic coastal mangrove lucinid species. The ISME Journal 2019; 13: 902–920.

Livingstone DR. Origins and Evolution of Pathways of Anaerobic Metabolism in the Animal Kingdom1. American Zoologist 1991; 31: 522–534.

Lobo-da-Cunha A. Structure and function of the digestive system in molluscs. Cell and Tissue Research 2019; 377: 475–503.

Lockwood BL, Connor KM, Gracey AY. The environmentally tuned transcriptomes of Mytilus mussels. J Exp Biol 2015; 218: 1822–33.

Loenen WAM. S-Adenosylmethionine: jack of all trades and master of everything? Biochemical Society Transactions 2006; 34: 330–333.

Love MI, Huber W, Anders S. Moderated estimation of fold change and dispersion for RNA-seq data with DESeq2. Genome Biology 2014; 15: 550.

Lumor L, Bock C, Mark FC, Ponsuksili S, Sokolova I. Effects of hypoxia–reoxygenation on the bioenergetics and oxidative stress in the isolated mitochondria of the king scallop, Pecten maximus. Journal of Experimental Biology 2025; 228: jeb249870.

Meng J, Wang T, Li B, Li L, Zhang G. Oxygen sensing and transcriptional regulation under hypoxia exposure in the mollusk Crassostrea gigas. Science of The Total Environment 2022; 853: 158557.

Milacic M, Beavers D, Conley P, Gong C, Gillespie M, Griss J, et al. The Reactome Pathway Knowledgebase 2024. Nucleic Acids Res 2024; 52: D672–d678.

Milacic M, Beavers D, Conley P, Gong C, Gillespie M, Griss J, et al. The Reactome Pathway Knowledgebase 2024. Nucleic Acids Research 2023; 52: D672–D678.

Mohanty RN, Clemens SC, Gupta AK. Dynamic shifts in the southern Benguela upwelling system since the latest Miocene. Earth and Planetary Science Letters 2024; 637: 118729.

Mohrholz V, Bartholomae CH, van der Plas AK, Lass HU. The seasonal variability of the northern Benguela undercurrent and its relation to the oxygen budget on the shelf. Continental Shelf Research 2008; 28: 424–441.

Montúfar-Romero M, Valenzuela-Muñoz V, Valenzuela-Miranda D, Gallardo-Escárate C. Hypoxia in the Blue Mussel Mytilus chilensis Induces a Transcriptome Shift Associated with Endoplasmic Reticulum Stress, Metabolism, and Immune Response. Genes (Basel) 2024; 15.

Morton JE. Molluscs: An Introduction to their Form and Function.. New York, USA: Harper Textbooks, 1960.

Nagaraj SH, Deshpande N, Gasser RB, Ranganathan S. ESTExplorer: an expressed sequence tag (EST) assembly and annotation platform. Nucleic Acids Research 2007; 35: W143–W147.

Nicholls DG. Mitochondrial proton leaks and uncoupling proteins. Biochim Biophys Acta Bioenerg 2021; 1862: 148428.

O’Farrell PH. Conserved responses to oxygen deprivation. J Clin Invest 2001; 107: 671–4.

Ortmann C, Grieshaber MK. Energy metabolism and valve closure behaviour in the Asian clam Corbicula fluminea. J Exp Biol 2003; 206: 4167–78.

Osvatic JT, Yuen B, Kunert M, Wilkins L, Hausmann B, Girguis P, et al. Gene loss and symbiont switching during adaptation to the deep sea in a globally distributed symbiosis. The ISME Journal 2023; 17: 453–466.

Pang Z, Lu Y, Zhou G, Hui F, Xu L, Viau C, et al. MetaboAnalyst 6.0: towards a unified platform for metabolomics data processing, analysis and interpretation. Nucleic Acids Research 2024; 52: W398–W406.

Pappalardo AM, Ferrito V, Biscotti MA, Canapa A, Capriglione T. Transposable Elements and Stress in Vertebrates: An Overview. Int J Mol Sci 2021; 22.

Park S, Cho JH, Kim JH, Park M, Park S, Kim SY, et al. Hypoxia stabilizes SETDB1 to maintain genome stability. Nucleic Acids Res 2023; 51: 11178–11196.

Pörtner HO, Heisler N, Grieshaber MK. Anaerobiosis and acid-base status in marine invertebrates: a theoretical analysis of proton generation by anaerobic metabolism. Journal of Comparative Physiology B 1984; 155: 1–12.

Rothfels K, Milacic M, Matthews L, Haw R, Sevilla C, Gillespie M, et al. Using the Reactome Database. Current Protocols 2023; 3: e722.

Schmidt-Rohr K. Oxygen Is the High-Energy Molecule Powering Complex Multicellular Life: Fundamental Corrections to Traditional Bioenergetics. ACS Omega 2020; 5: 2221–2233.

Scholz F. Identifying oxygen minimum zone-type biogeochemical cycling in Earth history using inorganic geochemical proxies. Earth-Science Reviews 2018; 184: 29–45.

Schreurs V, Aarts MJ, Ijssennagger N, Hermans J, Hendriks WH. Energetic and metabolic consequences of aerobic and anaerobic ATP-production. Agro Food Industry Hi-Tech 18 (2007) 5 2007; 18.

Semenza GL. Hydroxylation of HIF-1: Oxygen Sensing at the Molecular Level. Physiology 2004; 19: 176–182.

Shin J-S, Song C-u, Choi H, Yang SH, Kwon KK, Eyun S-i, et al. The Complete Mitochondrial Genome of the Chemosymbiotic Lucinid Bivalve Pillucina pisidium (Dunker, 1860) Occurring in Seagrass Zostera marina Bed in a Lagoon in Jeju Island, Korea. Journal of Marine Science and Engineering 2024; 12: 847.

Sokolov EP, Adzigbli L, Markert S, Bundgaard A, Fago A, Becher D, et al. Intrinsic Mechanisms Underlying Hypoxia-Tolerant Mitochondrial Phenotype During Hypoxia-Reoxygenation Stress in a Marine Facultative Anaerobe, the Blue Mussel Mytilus edulis. Frontiers in Marine Science 2021; 8.

Sokolova I. Bioenergetics in environmental adaptation and stress tolerance of aquatic ectotherms: linking physiology and ecology in a multi-stressor landscape. J Exp Biol 2021; 224.

Steffen JBM, Haider F, Sokolov EP, Bock C, Sokolova IM. Mitochondrial capacity and reactive oxygen species production during hypoxia and reoxygenation in the ocean quahog, Arctica islandica. Journal of Experimental Biology 2021; 224.

Steffen JBM, Sokolov EP, Bock C, Sokolova IM. Combined effects of salinity and intermittent hypoxia on mitochondrial capacity and reactive oxygen species efflux in the Pacific oyster, Crassostrea gigas. Journal of Experimental Biology 2023; 226.

Steinbrenner H, Speckmann B, Klotz L-O. Selenoproteins: Antioxidant selenoenzymes and beyond. Archives of Biochemistry and Biophysics 2016; 595: 113–119.

Taylor J, Glover E, Smith L, Ikebe C, Williams S. New molecular phylogeny of Lucinidae: Increased taxon base with focus on tropical Western Atlantic species (Mollusca: Bivalvia). Zootaxa 2016; 4196: 381.

Taylor JD, Glover EA, Smith L, Dyal P, Williams ST. Molecular phylogeny and classification of the chemosymbiotic bivalve family Lucinidae (Mollusca: Bivalvia). Zoological Journal of the Linnean Society 2011; 163: 15–49.

Tielens AG, Rotte C, van Hellemond JJ, Martin W. Mitochondria as we don’t know them. Trends Biochem Sci 2002; 27: 564–72.

Van den Thillart G, Verbeek R. Anoxia-Induced Oxygen Debt of Goldfish (Carassius auratus L.). Physiological Zoology 1991; 64: 525–540.

Vismann B, Hagerman L. Recovery from hypoxia with and without sulfide in Saduria entomon: oxygen debt, reduced sulfur and anaerobic metabolites. Marine Ecology Progress Series 1996; 143: 131–139.

Webster KA. Evolution of the coordinate regulation of glycolytic enzyme genes by hypoxia. Journal of Experimental Biology 2003; 206: 2911–2922.

Wu F, Kong H, Xie L, Sokolova IM. Exposure to nanopollutants (nZnO) enhances the negative effects of hypoxia and delays recovery of the mussels’ immune system. Environ Pollut 2024a; 351: 124112.

Wu F, Sokolov EP, Timm S, Sokolova IM. Synergistic impacts of nanopollutants (nZnO) and hypoxia on bioenergetics and metabolic homeostasis in a marine bivalve Mytilus edulis. Environmental Science: Nano 2024b.

Xie Y, Su N, Yang J, Tan Q, Huang S, Jin M, et al. FGF/FGFR signaling in health and disease. Signal Transduction and Targeted Therapy 2020; 5: 181.

Yang C, Wu H, Chen J, Liao Y, Mkuye R, Deng Y, et al. Integrated transcriptomic and metabolomic analysis reveals the response of pearl oyster (Pinctada fucata martensii) to long-term hypoxia. Marine Environmental Research 2023; 191: 106133.

Yuen B, Polzin J, Petersen JM. Organ transcriptomes of the lucinid clam Loripes orbiculatus (Poli, 1791) provide insights into their specialised roles in the biology of a chemosymbiotic bivalve. BMC Genomics 2019; 20: 820.

Zettler M, Bochert R, Pollehne F. Macrozoobenthic biodiversity patterns in the northern province of the Benguela upwelling system. African Journal of Marine Science 2013; 35: 283–290.

