## SupplementaryMaterials for "Unraveling Hypoxia Tolerance: Transcriptomic and Metabolic Insights from *Lucinoma capensis* in an Oxygen Minimum Zone"

**Supplementary Table 1.** List of the functional categories used for classification of eukaryotic and prokaryotic DEGs in the manually curated database of *L. capensis*.

| **Eukaryotic DEGs** | **Prokaryotic DEGs** |
|---|---|
| Acid-base regulation | Amino acid metabolism |
| Adhesion | Antibiotic metabolism |
| Amino acid metabolism | Apoptosis |
| Angiopoesis | ATP synthase |
| Apoptosis | Carbohydrate metabolism |
| ATP synthase (incl. assembly) | Cell division |
| Autophagy | Cell motility |
| Blood clotting | Cell wall |
| Ca metabolism | Chaperones |
| Carbohydrate metabolism | $CO_2$ fixation |
| Cell division & differentiation | Cofactor biosynthesis & metabolism |
| Chaperones | Deacetylase |
| Ciliary activity | Detoxification |
| Cofactor metabolism & biosynthesis | DNA maintenance |
| Cytoskeleton | ETS complexes (incl. assembly) |
| Detoxification | Glycolysis |
| Development | Hydrocarbon & alcohol metabolism |
| Digestion | Ion transport |
| DNA maintenance | Lipid metabolism |
| Endocrine function | Lipopolysaccharide biosynthesis |
| Endocytosis | Membrane transport |
| ETS complexes (incl. assembly) | Metal homeostasis |
| Extracellular matrix | Mobile genetic elements |
| Glycoprotein metabolism | Nitrogen fixation |
| Glycosphingolipid biosynthesis | NO metabolism |
| GSH metabolism | Nucleotide metabolism |
| Histone modification | Oxidoreductases |
| Immunity | Peptidoglycan synthesis |
| Inflammation | Phosphate metabolism and transport |
| Ion transport | Protein synthesis |
| Iron homeostasis | Protein transport |
| Ketone metabolism | Proteolysis & protein repair |
| Lipid metabolism | Recombination |
| Locomotion | Redox homeostasis |
| Membrane and substrate transport | RNA degradation |
| Membrane function | Signaling |
| miRNA transport | Sulfur metabolism |
| Mobile genetic elements | Transcription regulation |
| Neural function | Tricarboxylic acid cycle (TCA) |
| NO metabolism | Urea metabolism |
| Nuclear transport | Virulence factor |

| |
|---|
| Nucleotide metabolism |
| Oogenesis |
| Organelle biogenesis |
| Osmoregulation |
| Oxidoreductases |
| Phosphate metabolism and transport |
| Post-translational protein modifications |
| Protein synthesis |
| Protein transport |
| Proteolysis |
| Redox homeostasis |
| RNA metabolism |
| Secretion |
| Signaling |
| Spermatogenesis |
| Sphingolipid metabolism |
| Steroid metabolism & biosynthesis |
| Stress response |
| Sulfur metabolism |
| Transcription regulation |
| Tricarboxylic acid cycle (TCA) |
| Urea metabolism |
| Vision |

**Supplementary Table 2.** Relative frequency of functional gene categories (as listed in Supplementary Table 1) among differentially expressed eukaryotic genes (DEGs) identified in the gill tissues of *L. capensis* exposed to varying oxygen conditions.

The table displays the percentage of total DEGs for each category and the cumulative percentage (Cum. %) of categories, ranked in descending order of DEG count.

| **A. Downregulated in hypoxia vs. normoxia (819 DEGs)** | | | | **B. Upregulated in hypoxia vs. normoxia (236 DEGs)** | | | |
|---|---|---|---|---|---|---|---|
| **Functional group** | **# DEGs** | **% of total** | **Cum. %** | **Functional group** | **# DEGs** | **% of total** | **Cum. %** |
| Protein synthesis | 150 | 18.3 | 18.3 | Proteolysis | 25 | 10.6 | 10.6 |
| Proteolysis | 62 | 7.6 | 25.9 | Transcription regulation | 18 | 7.6 | 18.2 |
| Mobile genetic elements (transposition) | 58 | 7.1 | 33.0 | Immunity | 17 | 7.2 | 25.4 |
| Immunity | 52 | 6.3 | 39.3 | Cytoskeleton | 17 | 7.2 | 32.6 |
| Signaling | 47 | 5.7 | 45.1 | Carbohydrate metabolism | 15 | 6.4 | 39.0 |
| Transcription regulation | 40 | 4.9 | 49.9 | DNA maintenance | 13 | 5.5 | 44.5 |
| ETS complexes (incl. assembly) | 32 | 3.9 | 53.8 | ECM | 13 | 5.5 | 50.0 |
| Detoxification | 23 | 2.8 | 56.7 | Protein synthesis | 12 | 5.1 | 55.1 |
| Neural function | 23 | 2.8 | 59.5 | Chaperone | 10 | 4.2 | 59.3 |
| Carbohydrate metabolism | 22 | 2.7 | 62.1 | Signaling | 8 | 3.4 | 62.7 |
| DNA maintenance | 21 | 2.6 | 64.7 | Adhesion | 7 | 3.0 | 65.7 |
| AA metabolism | 21 | 2.6 | 67.3 | Mobile genetic elements (transposition) | 7 | 3.0 | 68.6 |
| Lipid metabolism | 19 | 2.3 | 69.6 | Redox homeostasis | 7 | 3.0 | 71.6 |
| RNA metabolism | 16 | 2.0 | 71.6 | ETS complexes (incl. assembly) | 6 | 2.5 | 74.2 |
| Cytoskeleton | 15 | 1.8 | 73.4 | Cell division & differentiation | 5 | 2.1 | 76.3 |
| ECM | 14 | 1.7 | 75.1 | Nucleotide metabolism | 5 | 2.1 | 78.4 |
| Cell division & differentiation | 13 | 1.6 | 76.7 | Endocytosis | 4 | 1.7 | 80.1 |
| Redox homeostasis | 13 | 1.6 | 78.3 | Locomotion | 4 | 1.7 | 81.8 |

| | | | | | | | |
|---|---|---|---|---|---|---|---|
| Chaperone | 12 | 1.5 | 79.7 | Membrane and substrate transport | 4 | 1.7 | 83.5 |
| Apoptosis | 12 | 1.5 | 81.2 | AA metabolism | 3 | 1.3 | 84.7 |
| Membrane and substrate transport | 12 | 1.5 | 82.7 | Ca metabolism | 3 | 1.3 | 86.0 |
| Cofactor metabolism & biosynthesis | 10 | 1.2 | 83.9 | Detoxification | 3 | 1.3 | 87.3 |
| Adhesion | 9 | 1.1 | 85.0 | Inflammation | 3 | 1.3 | 88.6 |
| Ca metabolism | 9 | 1.1 | 86.1 | Spermatogenesis | 3 | 1.3 | 89.8 |
| Nucleotide metabolism | 9 | 1.1 | 87.2 | Neural function | 3 | 1.3 | 91.1 |
| Locomotion | 8 | 1.0 | 88.2 | RNA metabolism | 3 | 1.3 | 92.4 |
| Histone modification | 7 | 0.9 | 89.0 | Development | 2 | 0.8 | 93.2 |
| Sulfur metabolism | 7 | 0.9 | 89.9 | Ciliary activity | 2 | 0.8 | 94.1 |
| Protein transport | 6 | 0.7 | 90.6 | Ion transport | 2 | 0.8 | 94.9 |
| Ciliary activity | 5 | 0.6 | 91.2 | Cofactor metabolism & biosynthesis | 2 | 0.8 | 95.8 |
| Membrane function | 5 | 0.6 | 91.8 | Sulfur metabolism | 2 | 0.8 | 96.6 |
| Spermatogenesis | 5 | 0.6 | 92.4 | Apoptosis | 1 | 0.4 | 97.0 |
| Sphingolipid metabolism | 5 | 0.6 | 93.0 | Glycoprotein metabolism | 1 | 0.4 | 97.5 |
| ATP synthase (incl. assembly) | 5 | 0.6 | 93.7 | Glycosphingolipid biosynthesis | 1 | 0.4 | 97.9 |
| Inflammation | 4 | 0.5 | 94.1 | Lipid metabolism | 1 | 0.4 | 98.3 |
| Ion transport | 4 | 0.5 | 94.6 | Membrane function | 1 | 0.4 | 98.7 |
| PTM | 4 | 0.5 | 95.1 | Nuclear transport | 1 | 0.4 | 99.2 |
| Stress response | 4 | 0.5 | 95.6 | Oogenesis | 1 | 0.4 | 99.6 |
| Steroid metabolism & biosynthesis | 4 | 0.5 | 96.1 | Oxidoreductases | 1 | 0.4 | 100.0 |
| Phosphate metabolism and transport | 3 | 0.4 | 96.5 | | | | |
| Acid-base regulation | 3 | 0.4 | 96.8 | | | | |
| Nuclear transport | 3 | 0.4 | 97.2 | | | | |
| Blood clotting | 2 | 0.2 | 97.4 | | | | |
| Development | 2 | 0.2 | 97.7 | | | | |
| Endocytosis | 2 | 0.2 | 97.9 | | | | |

| | | | | | | | |
|---|---|---|---|---|---|---|---|
| Angiopoesis | 2 | 0.2 | 98.2 | | | | |
| Endocrine function | 2 | 0.2 | 98.4 | | | | |
| Urea metabolism | 2 | 0.2 | 98.7 | | | | |
| Iron homeostasis | 2 | 0.2 | 98.9 | | | | |
| NO metabolism | 1 | 0.1 | 99.0 | | | | |
| Autophagy | 1 | 0.1 | 99.1 | | | | |
| Digestion | 1 | 0.1 | 99.3 | | | | |
| Glycoprotein metabolism | 1 | 0.1 | 99.4 | | | | |
| Ketone metabolism | 1 | 0.1 | 99.5 | | | | |
| miRNA transport | 1 | 0.1 | 99.6 | | | | |
| Vision? | 1 | 0.1 | 99.8 | | | | |
| Organelle biogenesis | 1 | 0.1 | 99.9 | | | | |
| TCA | 1 | 0.1 | 100.0 | | | | |
| **C. Downregulated in 1 h recovery vs. normoxia (409 DEGs)** | | | | **D. Upregulated in 1 h recovery vs. normoxia (289 DEGs)** | | | |
| **Functional group** | **# DEGs** | **% of total** | **Cum. %** | **Functional group** | **# DEGs** | **% of total** | **Cum. %** |
| Proteolysis incl. ubiquitin-proteasome pathway | 49 | 12.0 | 12.0 | Proteolysis incl. ubiquitin-proteasome pathway | 27 | 9.3 | 9.3 |
| Mobile genetic elements (transposition) | 43 | 10.5 | 22.5 | ECM | 19 | 6.6 | 15.9 |
| Immunity | 30 | 7.3 | 29.8 | Mobile genetic elements (transposition) | 17 | 5.9 | 21.8 |
| Transcription regulation & transcription factors | 25 | 6.1 | 35.9 | Immunity | 17 | 5.9 | 27.7 |
| Protein synthesis | 23 | 5.6 | 41.6 | Transcription regulation & transcription factors | 17 | 5.9 | 33.6 |
| Signaling | 19 | 4.6 | 46.2 | Signaling | 16 | 5.5 | 39.1 |
| Apoptosis | 16 | 3.9 | 50.1 | Cytoskeleton | 14 | 4.8 | 43.9 |
| DNA maintenance | 15 | 3.7 | 53.8 | Protein synthesis | 12 | 4.2 | 48.1 |

| | | | | | | | |
|---|---|---|---|---|---|---|---|
| Detoxification | 12 | 2.9 | 56.7 | Lipid metabolism | 11 | 3.8 | 51.9 |
| Lipid metabolism | 12 | 2.9 | 59.7 | DNA maintenance | 10 | 3.5 | 55.4 |
| Cytoskeleton | 10 | 2.4 | 62.1 | Adhesion | 9 | 3.1 | 58.5 |
| Neural function | 10 | 2.4 | 64.5 | Chaperone | 8 | 2.8 | 61.2 |
| Membrane and substrate transport | 10 | 2.4 | 67.0 | Neural function | 9 | 3.1 | 64.4 |
| Cell division & differentiation | 9 | 2.2 | 69.2 | Carbohydrate metabolism | 7 | 2.4 | 66.8 |
| Carbohydrate metabolism | 9 | 2.2 | 71.4 | Cell division & differentiation | 7 | 2.4 | 69.2 |
| Nucleotide metabolism | 8 | 2.0 | 73.3 | RNA metabolism | 7 | 2.4 | 71.6 |
| Redox homeostasis | 8 | 2.0 | 75.3 | AA metabolism | 6 | 2.1 | 73.7 |
| AA metabolism | 7 | 1.7 | 77.0 | Locomotion | 6 | 2.1 | 75.8 |
| RNA metabolism | 7 | 1.7 | 78.7 | Detoxification | 5 | 1.7 | 77.5 |
| Chaperone | 6 | 1.5 | 80.2 | Nucleotide metabolism | 5 | 1.7 | 79.2 |
| ECM | 6 | 1.5 | 81.7 | Redox homeostasis | 4 | 1.4 | 80.6 |
| Histone modification | 6 | 1.5 | 83.1 | Inflammation | 4 | 1.4 | 82.0 |
| Adhesion | 5 | 1.2 | 84.4 | Development | 4 | 1.4 | 83.4 |
| Oxidoreductases | 5 | 1.2 | 85.6 | Membrane and substrate transport | 4 | 1.4 | 84.8 |
| Inflammation | 4 | 1.0 | 86.6 | Ca metabolism | 3 | 1.0 | 85.8 |
| Locomotion | 4 | 1.0 | 87.5 | Spermatogenesis | 3 | 1.0 | 86.9 |
| Cofactor metabolism & biosynthesis | 4 | 1.0 | 88.5 | Nuclear transport | 3 | 1.0 | 87.9 |
| ETS complexes (incl. assembly) | 4 | 1.0 | 89.5 | Ciliary activity | 3 | 1.0 | 88.9 |
| Ca metabolism | 3 | 0.7 | 90.2 | Cofactor metabolism & biosynthesis | 3 | 1.0 | 90.0 |
| Ion transport | 3 | 0.7 | 91.0 | ETS complexes (incl. assembly) | 3 | 1.0 | 91.0 |
| Membrane function | 3 | 0.7 | 91.7 | Endocytosis | 2 | 0.7 | 91.7 |
| Nuclear transport | 3 | 0.7 | 92.4 | Glycoprotein metabolism | 2 | 0.7 | 92.4 |
| Sulfur metabolism | 3 | 0.7 | 93.2 | Histone modification | 2 | 0.7 | 93.1 |
| Endocytosis | 3 | 0.7 | 93.9 | Phosphate metabolism and transport | 2 | 0.7 | 93.8 |
| Steroid metabolism & biosynthesis | 3 | 0.7 | 94.6 | Protein transport | 2 | 0.7 | 94.5 |

| | | | | | | | |
|---|---|---|---|---|---|---|---|
| Iron homeostasis | 3 | 0.7 | 95.4 | Stress response | 2 | 0.7 | 95.2 |
| Acid-base regulation | 2 | 0.5 | 95.8 | Sulfur metabolism | 2 | 0.7 | 95.8 |
| Protein transport | 2 | 0.5 | 96.3 | Osmoregulation | 2 | 0.7 | 96.5 |
| Stress response | 2 | 0.5 | 96.8 | Apoptosis | 2 | 0.7 | 97.2 |
| Autophagy | 1 | 0.2 | 97.1 | Oxidoreductases | 2 | 0.7 | 97.9 |
| Ciliary activity | 1 | 0.2 | 97.3 | Autophagy | 1 | 0.3 | 98.3 |
| Glycoprotein metabolism | 1 | 0.2 | 97.6 | Ion transport | 1 | 0.3 | 98.6 |
| Ketone metabolism | 1 | 0.2 | 97.8 | Iron homeostasis | 1 | 0.3 | 99.0 |
| Oogenesis | 1 | 0.2 | 98.0 | PTM | 1 | 0.3 | 99.3 |
| Organelle biogenesis | 1 | 0.2 | 98.3 | Sphingolipid metabolism | 1 | 0.3 | 99.7 |
| Osmoregulation | 1 | 0.2 | 98.5 | Steroid metabolism & biosynthesis | 1 | 0.3 | 100.0 |
| Phosphate metabolism and transport | 1 | 0.2 | 98.8 | | | | |
| Spermatogenesis | 1 | 0.2 | 99.0 | | | | |
| Sphingolipid metabolism | 1 | 0.2 | 99.3 | | | | |
| Urea metabolism | 1 | 0.2 | 99.5 | | | | |
| ATP synthase (incl. assembly) | 1 | 0.2 | 99.8 | | | | |
| TCA | 1 | 0.2 | 100.0 | | | | |
| **E. Downregulated in 24 h recovery vs. normoxia (261 DEGs)** | | | | **F. Upregulated in 24 h recovery vs. normoxia (328 DEGs)** | | | |
| **Functional group** | **# DEGs** | **% of total** | **Cum. %** | **Functional group** | **# DEGs** | **% of total** | **Cum. %** |
| Immunity | 29 | 11.1 | 11.1 | Proteolysis incl. ubiquitin-proteasome pathway | 27 | 8.2 | 8.2 |
| Proteolysis incl. ubiquitin-proteasome pathway | 23 | 8.8 | 19.9 | Transcription regulation & transcription factors | 25 | 7.6 | 15.9 |
| Protein synthesis | 20 | 7.7 | 27.6 | DNA maintenance | 23 | 7.0 | 22.9 |
| Mobile genetic elements (transposition) | 18 | 6.9 | 34.5 | Mobile genetic elements (transposition) | 20 | 6.1 | 29.0 |

| | | | | | | | |
|---|---|---|---|---|---|---|---|
| Chaperone | 11 | 4.2 | 38.7 | Signaling | 17 | 5.2 | 34.1 |
| Transcription regulation & transcription factors | 11 | 4.2 | 42.9 | Locomotion | 14 | 4.3 | 38.4 |
| Neural function | 10 | 3.8 | 46.7 | Cytoskeleton | 12 | 3.7 | 42.1 |
| Lipid metabolism | 8 | 3.1 | 49.8 | Protein synthesis | 12 | 3.7 | 45.7 |
| ECM | 7 | 2.7 | 52.5 | RNA metabolism | 12 | 3.7 | 49.4 |
| Detoxification | 7 | 2.7 | 55.2 | Immunity | 11 | 3.4 | 52.7 |
| Carbohydrate metabolism | 7 | 2.7 | 57.9 | Adhesion | 10 | 3.0 | 55.8 |
| Signaling | 6 | 2.3 | 60.2 | Chaperone | 9 | 2.7 | 58.5 |
| AA metabolism | 6 | 2.3 | 62.5 | ECM | 9 | 2.7 | 61.3 |
| Cell division & differentiation | 6 | 2.3 | 64.8 | Cell division & differentiation | 9 | 2.7 | 64.0 |
| Locomotion | 6 | 2.3 | 67.0 | Ciliary activity | 9 | 2.7 | 66.8 |
| DNA maintenance | 6 | 2.3 | 69.3 | Neural function | 9 | 2.7 | 69.5 |
| RNA metabolism | 6 | 2.3 | 71.6 | PTM | 8 | 2.4 | 72.0 |
| Histone modification | 5 | 1.9 | 73.6 | Histone modification | 7 | 2.1 | 74.1 |
| Ion transport | 5 | 1.9 | 75.5 | Spermatogenesis | 7 | 2.1 | 76.2 |
| Endocytosis | 5 | 1.9 | 77.4 | Carbohydrate metabolism | 7 | 2.1 | 78.4 |
| Apoptosis | 4 | 1.5 | 78.9 | Stress response | 6 | 1.8 | 80.2 |
| Cytoskeleton | 4 | 1.5 | 80.5 | Ca metabolism | 6 | 1.8 | 82.0 |
| GSH metabolism | 3 | 1.1 | 81.6 | Endocytosis | 5 | 1.5 | 83.5 |
| Oogenesis | 3 | 1.1 | 82.8 | ETS complexes (incl. assembly) | 5 | 1.5 | 85.1 |
| PTM | 3 | 1.1 | 83.9 | Development | 4 | 1.2 | 86.3 |
| Iron homeostasis | 3 | 1.1 | 85.1 | Lipid metabolism | 4 | 1.2 | 87.5 |
| Ciliary activity | 3 | 1.1 | 86.2 | Nucleotide metabolism | 4 | 1.2 | 88.7 |
| ETS complexes (incl. assembly) | 3 | 1.1 | 87.4 | Apoptosis | 3 | 0.9 | 89.6 |
| Acid-base regulation | 2 | 0.8 | 88.1 | Detoxification | 3 | 0.9 | 90.5 |
| Autophagy | 2 | 0.8 | 88.9 | Ion transport | 3 | 0.9 | 91.5 |
| Development | 2 | 0.8 | 89.7 | Membrane and substrate transport | 3 | 0.9 | 92.4 |
| Ketone metabolism | 2 | 0.8 | 90.4 | Nuclear transport | 3 | 0.9 | 93.3 |

| | | | | | | | |
|---|---|---|---|---|---|---|---|
| Oxidoreductases | 2 | 0.8 | 91.2 | Redox homeostasis | 3 | 0.9 | 94.2 |
| Protein transport | 2 | 0.8 | 92.0 | Membrane function | 2 | 0.6 | 94.8 |
| PTM | 2 | 0.8 | 92.7 | Organelle biogenesis | 2 | 0.6 | 95.4 |
| Spermatogenesis | 2 | 0.8 | 93.5 | Secretion | 2 | 0.6 | 96.0 |
| Membrane and substrate transport | 2 | 0.8 | 94.3 | Membrane and substrate transport | 2 | 0.6 | 96.6 |
| Sulfur metabolism | 2 | 0.8 | 95.0 | ATP synthase (incl. assembly) | 2 | 0.6 | 97.3 |
| Nucleotide metabolism | 2 | 0.8 | 95.8 | AA metabolism | 2 | 0.6 | 97.9 |
| Adhesion | 1 | 0.4 | 96.2 | Cofactor metabolism & biosynthesis | 2 | 0.6 | 98.5 |
| Ca metabolism | 1 | 0.4 | 96.6 | Oxidoreductases | 2 | 0.6 | 99.1 |
| Cofactor metabolism & biosynthesis | 1 | 0.4 | 96.9 | Glycoprotein metabolism | 1 | 0.3 | 99.4 |
| Endocrine function | 1 | 0.4 | 97.3 | Inflammation | 1 | 0.3 | 99.7 |
| Nuclear transport | 1 | 0.4 | 97.7 | Protein transport | 1 | 0.3 | 100.0 |
| Phosphate metabolism and transport | 1 | 0.4 | 98.1 | | | | |
| Secretion | 1 | 0.4 | 98.5 | | | | |
| Sphingolipid metabolism | 1 | 0.4 | 98.9 | | | | |
| Steroid metabolism & biosynthesis | 1 | 0.4 | 99.2 | | | | |
| Stress response | 1 | 0.4 | 99.6 | | | | |
| Urea metabolism | 1 | 0.4 | 100.0 | | | | |

**Supplementary Table 3.** Relative frequency of functional gene categories (as listed in Supplementary Table 1) among differentially expressed prokaryotic genes (DEGs) identified in the gill tissues of *L. capensis* exposed to varying oxygen conditions.

The table displays the percentage of total DEGs for each category and the cumulative percentage (Cum. %) of categories, ranked in descending order of DEG count.

| A. Downregulated in anoxia vs. normoxia (40) | | | | B. Upregulated in anoxia vs. normoxia (139) | | | |
|---|---|---|---|---|---|---|---|
| **Functional group** | **# DEGs** | **% of total** | **Cum.%** | **Functional group** | **# DEGs** | **% of total** | **Cum.%** |
| Antibiotic metabolism | 4 | 10 | 10 | Sulfur metabolism | 28 | 20.1 | 20.1 |
| Amino acid metabolism | 4 | 10 | 20 | CO2 fixation | 18 | 12.9 | 33.1 |
| Proteolysis & protein repair | 3 | 7.5 | 27.5 | ETS | 16 | 11.5 | 44.6 |
| Chaperone | 2 | 5 | 32.5 | Protein synthesis and transport | 15 | 10.8 | 55.4 |
| CO2 fixation | 2 | 5 | 37.5 | Amino acid metabolism | 14 | 10.1 | 65.5 |
| Phosphate metabolism | 2 | 5 | 42.5 | Nucleotide metabolism | 6 | 4.3 | 69.8 |
| Mobile genetic elements | 2 | 5 | 47.5 | NO metabolism | 4 | 2.9 | 72.7 |
| Hydrocarbon & alcohol metabolism | 2 | 5 | 52.5 | Proteolysis & protein repair | 4 | 2.9 | 75.5 |
| Cofactor biosynthesis & metabolism | 2 | 5 | 57.5 | Membrane transport | 4 | 2.9 | 78.4 |
| Transcription regulation | 2 | 5 | 62.5 | ATP synthase | 3 | 2.2 | 80.6 |
| Metal homeostasis | 2 | 5 | 67.5 | Chaperone | 3 | 2.2 | 82.7 |
| Lipopolysaccharide biosynthesis | 2 | 5 | 72.5 | Peptidoglycan synthesis | 3 | 2.2 | 84.9 |
| DNA maintenance | 1 | 2.5 | 75 | Redox homeostasis | 3 | 2.2 | 87.1 |
| ETS | 1 | 2.5 | 77.5 | Cell division | 2 | 1.4 | 88.5 |
| Nitrogen fixation | 1 | 2.5 | 80 | Lipopolysaccharide biosynthesis | 2 | 1.4 | 89.9 |
| NO metabolism | 1 | 2.5 | 82.5 | Phosphate metabolism | 2 | 1.4 | 91.4 |
| Oxidoreductases | 1 | 2.5 | 85 | Signaling | 2 | 1.4 | 92.8 |
| Peptidoglycan synthesis | 1 | 2.5 | 87.5 | Tricarboxylic acid cycle | 2 | 1.4 | 94.2 |
| Protein synthesis and transport | 1 | 2.5 | 90 | DNA maintenance | 2 | 1.4 | 95.7 |
| Recombination | 1 | 2.5 | 92.5 | Transcription regulation | 2 | 1.4 | 97.1 |
| Signaling | 1 | 2.5 | 95 | Lipid metabolism | 1 | 0.7 | 97.8 |

| | | | | | | | |
|---|---|---|---|---|---|---|---|
| Tricarboxylic acid cycle | 1 | 2.5 | 97.5 | RNA degradation | 1 | 0.7 | 98.6 |
| Virulence factor | 1 | 2.5 | 100 | Mobile genetic elements (transposition) | 1 | 0.7 | 99.3 |
| | | | | Virulence factor | 1 | 0.7 | 100.0 |
| **C. Downregulated in 1 h recovery vs. normoxia (88)** | | | | **D. Upregulated in 1 h recovery vs. normoxia (52)** | | | |
| **Functional group** | **# DEGs** | **% of total** | **Cum. %** | **Functional group** | **# DEGs** | **% of total** | **Cum. %** |
| Amino acid metabolism | 11 | 12.5 | 12.5 | CO2 fixation | 8 | 15.4 | 15.4 |
| Protein synthesis and transport | 8 | 9.1 | 21.6 | NO metabolism | 6 | 11.5 | 26.9 |
| Mobile genetic elements (transposition) | 5 | 5.7 | 27.3 | Sulfur metabolism | 5 | 9.6 | 36.5 |
| Protein transport | 4 | 4.5 | 31.8 | Transcription regulation | 6 | 11.5 | 48.1 |
| Antibiotic metabolism | 3 | 3.4 | 35.2 | Tricarboxylic acid cycle | 3 | 5.8 | 53.8 |
| Chaperone | 3 | 3.4 | 38.6 | Amino acid metabolism | 3 | 5.8 | 59.6 |
| DNA maintenance | 3 | 3.4 | 42.0 | Antibiotic metabolism | 2 | 3.8 | 63.5 |
| Proteolysis & protein repair | 3 | 3.4 | 45.5 | Membrane transport | 2 | 3.8 | 67.3 |
| Redox homeostasis | 3 | 3.4 | 48.9 | Protein synthesis and transport | 2 | 3.8 | 71.2 |
| Tricarboxylic acid cycle | 3 | 3.4 | 52.3 | Glycolysis | 1 | 1.9 | 73.1 |
| Transcription regulation | 3 | 3.4 | 55.7 | ATP synthase | 1 | 1.9 | 75.0 |
| Cofactor biosynthesis & metabolism | 3 | 3.4 | 59.1 | Chaperone | 1 | 1.9 | 76.9 |
| Carbohydrate metabolism | 2 | 2.3 | 61.4 | Detoxification | 1 | 1.9 | 78.8 |
| Cell wall | 2 | 2.3 | 63.6 | DNA maintenance | 1 | 1.9 | 80.8 |
| CO2 fixation | 2 | 2.3 | 65.9 | ETS | 1 | 1.9 | 82.7 |
| Detoxification | 2 | 2.3 | 68.2 | Ion transport | 1 | 1.9 | 84.6 |
| ETS | 2 | 2.3 | 70.5 | Lipopolysaccharide biosynthesis | 1 | 1.9 | 86.5 |
| Glycolysis | 2 | 2.3 | 72.7 | Nucleotide metabolism | 2 | 3.8 | 90.4 |
| Lipid metabolism | 2 | 2.3 | 75.0 | Oxidoreductases | 1 | 1.9 | 92.3 |
| Membrane transport | 2 | 2.3 | 77.3 | Phosphate metabolism | 1 | 1.9 | 94.2 |
| Metal homeostasis | 2 | 2.3 | 79.5 | Proteolysis & protein repair | 1 | 1.9 | 96.2 |
| Oxidoreductases | 2 | 2.3 | 81.8 | Redox homeostasis | 1 | 1.9 | 98.1 |

| | | | | | | | |
|---|---|---|---|---|---|---|---|
| Signaling | 2 | 2.3 | 84.1 | Virulence factor | 1 | 1.9 | 100.0 |
| Nucleotide metabolism | 2 | 2.3 | 86.4 | | | | |
| Transcription regulation | 2 | 2.3 | 88.6 | | | | |
| Hydrocarbon and alcohol metabolism | 1 | 1.1 | 89.8 | | | | |
| Apoptosis | 1 | 1.1 | 90.9 | | | | |
| Ion transport | 1 | 1.1 | 92.0 | | | | |
| Lipopolysaccharide biosynthesis | 1 | 1.1 | 93.2 | | | | |
| NO metabolism | 1 | 1.1 | 94.3 | | | | |
| Phosphate metabolism and transport | 1 | 1.1 | 95.5 | | | | |
| Recombination | 1 | 1.1 | 96.6 | | | | |
| Sulfur metabolism | 1 | 1.1 | 97.7 | | | | |
| Urea metabolism | 1 | 1.1 | 98.9 | | | | |
| Virulence factor | 1 | 1.1 | 100.0 | | | | |
| **E. Downregulated in 24 h recovery vs. normoxia (18)** | | | | **F. Upregulated in 24 h recovery vs. normoxia (84)** | | | |
| **Functional group** | **# DEGs** | **% of total** | **Cum. %** | **Functional group** | **# DEGs** | **% of total** | **Cum. %** |
| Transcription regulation | 2 | 11.1 | 11.1 | Protein synthesis and transport | 8 | 9.5 | 9.5 |
| Antibiotic metabolism | 2 | 11.1 | 22.2 | CO2 fixation | 7 | 8.3 | 17.9 |
| Tricarboxylic acid cycle | 2 | 11.1 | 33.3 | Transcription regulation | 7 | 8.3 | 26.2 |
| Virulence factor | 2 | 11.1 | 44.4 | Sulfur metabolism | 7 | 8.3 | 34.5 |
| Cell wall | 2 | 11.1 | 55.6 | ETS | 6 | 7.1 | 41.7 |
| Chaperone | 1 | 5.6 | 61.1 | Chaperone | 5 | 6.0 | 47.6 |
| CO2 fixation | 1 | 5.6 | 66.7 | Amino acid metabolism | 5 | 6.0 | 53.6 |
| DNA maintenance | 1 | 5.6 | 72.2 | Nucleotide metabolism | 4 | 4.8 | 58.3 |
| Lipopolysaccharide biosynthesis | 1 | 5.6 | 77.8 | Cell wall | 4 | 4.8 | 63.1 |
| Phosphate metabolism | 1 | 5.6 | 83.3 | Lipid metabolism | 3 | 3.6 | 66.7 |
| Proteolysis & protein repair | 1 | 5.6 | 88.9 | Oxidoreductases | 3 | 3.6 | 70.2 |
| Sulfur metabolism | 1 | 5.6 | 94.4 | DNA maintenance | 3 | 3.6 | 73.8 |

| | | | | | | | |
|---|---|---|---|---|---|---|---|
| Mobile genetic elements (transposition) | 1 | 5.6 | 100.0 | Membrane transport | 3 | 3.6 | 77.4 |
| | | | | Glycolysis | 2 | 2.4 | 79.8 |
| | | | | NO metabolism | 2 | 2.4 | 82.1 |
| | | | | Redox homeostasis | 2 | 2.4 | 84.5 |
| | | | | Tricarboxylic acid cycle | 2 | 2.4 | 86.9 |
| | | | | Carbohydrate metabolism | 1 | 1.2 | 88.1 |
| | | | | Cell division | 1 | 1.2 | 89.3 |
| | | | | Cell motility | 1 | 1.2 | 90.5 |
| | | | | Cofactor biosynthesis & metabolism | 1 | 1.2 | 91.7 |
| | | | | Lipopolysaccharide biosynthesis | 1 | 1.2 | 92.9 |
| | | | | Phosphate metabolism | 1 | 1.2 | 94.0 |
| | | | | Proteolysis & protein repair | 1 | 1.2 | 95.2 |
| | | | | Signaling | 1 | 1.2 | 96.4 |
| | | | | Stress response | 1 | 1.2 | 97.6 |
| | | | | Mobile genetic elements (transposition) | 1 | 1.2 | 98.8 |
| | | | | Virulence factor | 1 | 1.2 | 100.0 |

**Supplementary Table 4.** Relative frequency of functional gene categories (as listed in Supplementary Table 1) among differentially expressed genes (DEGs) identified in the digestive gland tissues of *L. capensis* exposed to varying oxygen conditions.

The table displays the percentage of total DEGs for each category and the cumulative percentage (Cum. %) of categories, ranked in descending order of DEG count.

| **A. Downregulated in anoxia vs. normoxia (167)** | | | | **B. Upregulated in anoxia vs. normoxia (382)** | | | |
|---|---|---|---|---|---|---|---|
| **Functional group** | **# DEGs** | **% of total** | **Cum. %** | **Functional group** | **# DEGs** | **% of total** | Cum.% |
| Protein synthesis | 18 | 10.8 | 10.8 | Immunity | 62 | 16.2 | 16.2 |
| Mobile genetic elements (transposition) | 12 | 7.2 | 18.0 | Proteolysis incl. ubiquitin-proteasome pathway | 38 | 9.9 | 26.2 |
| Proteolysis incl. ubiquitin-proteasome pathway | 11 | 6.6 | 24.6 | ECM | 32 | 8.4 | 34.6 |
| Cell division & differentiation | 8 | 4.8 | 29.3 | Adhesion | 16 | 4.2 | 38.7 |
| Immunity | 7 | 4.2 | 33.5 | Protein synthesis | 16 | 4.2 | 42.9 |
| Signaling | 7 | 4.2 | 37.7 | Neural function | 14 | 3.7 | 46.6 |
| Transcription regulation & transcription factors | 7 | 4.2 | 41.9 | Lipid metabolism | 14 | 3.7 | 50.3 |
| Neural function | 7 | 4.2 | 46.1 | DNA maintenance | 13 | 3.4 | 53.7 |
| Apoptosis | 6 | 3.6 | 49.7 | Cytoskeleton | 11 | 2.9 | 56.5 |
| Ciliary activity | 6 | 3.6 | 53.3 | Signaling | 10 | 2.6 | 59.2 |
| RNA metabolism | 6 | 3.6 | 56.9 | Carbohydrate metabolism | 9 | 2.4 | 61.5 |
| Adhesion | 5 | 3.0 | 59.9 | Mobile genetic elements (transposition) | 9 | 2.4 | 63.9 |
| Cytoskeleton | 5 | 3.0 | 62.9 | Chaperone | 8 | 2.1 | 66.0 |
| ECM | 5 | 3.0 | 65.9 | Cofactor metabolism & biosynthesis | 8 | 2.1 | 68.1 |
| Cofactor metabolism & biosynthesis | 5 | 3.0 | 68.9 | Detoxification | 8 | 2.1 | 70.2 |
| Carbohydrate metabolism | 4 | 2.4 | 71.3 | Locomotion | 8 | 2.1 | 72.3 |
| Ion transport | 4 | 2.4 | 73.7 | Apoptosis | 7 | 1.8 | 74.1 |
| PTM | 4 | 2.4 | 76.0 | Transcription regulation & transcription factors | 7 | 1.8 | 75.9 |
| Membrane and substrate transport | 4 | 2.4 | 78.4 | RNA metabolism | 7 | 1.8 | 77.7 |

| | | | | | | | |
|---|---|---|---|---|---|---|---|
| Detoxification | 3 | 1.8 | 80.2 | AA metabolism | 6 | 1.6 | 79.3 |
| Redox homeostasis | 3 | 1.8 | 82.0 | Glycoprotein metabolism | 6 | 1.6 | 80.9 |
| Histone modification | 3 | 1.8 | 83.8 | Ion transport | 6 | 1.6 | 82.5 |
| Lipid metabolism | 3 | 1.8 | 85.6 | Oxidoreductases | 6 | 1.6 | 84.0 |
| Nucleotide metabolism | 3 | 1.8 | 87.4 | Redox homoestasis | 6 | 1.6 | 85.6 |
| Protein transport | 3 | 1.8 | 89.2 | Protein transport | 5 | 1.3 | 86.9 |
| AA metabolism | 3 | 1.8 | 91.0 | Cell division & differentiation | 5 | 1.3 | 88.2 |
| Nuclear transport | 2 | 1.2 | 92.2 | Ca metabolism | 4 | 1.0 | 89.3 |
| Sulfur metabolism | 2 | 1.2 | 93.4 | PTM | 4 | 1.0 | 90.3 |
| Urea metabolism | 2 | 1.2 | 94.6 | ETS complexes (incl. assembly) | 4 | 1.0 | 91.4 |
| DNA maintenance | 1 | 0.6 | 95.2 | Ciliary activity | 3 | 0.8 | 92.1 |
| Chaperone | 1 | 0.6 | 95.8 | Membrane function | 3 | 0.8 | 92.9 |
| Endocytosis | 1 | 0.6 | 96.4 | Stress response | 3 | 0.8 | 93.7 |
| Inflammation | 1 | 0.6 | 97.0 | Inflammation | 2 | 0.5 | 94.2 |
| Locomotion | 1 | 0.6 | 97.6 | Spermatogenesis | 2 | 0.5 | 94.8 |
| Oxidoreductases | 1 | 0.6 | 98.2 | Sphingolipid metabolism | 2 | 0.5 | 95.3 |
| Peptidoglycan synthesis | 1 | 0.6 | 98.8 | Sulfur metabolism | 2 | 0.5 | 95.8 |
| Spermatogenesis | 1 | 0.6 | 99.4 | Osmoregulation | 2 | 0.5 | 96.3 |
| ETS complexes (incl. assembly) | 1 | 0.6 | 100.0 | Angiogenesis | 1 | 0.3 | 96.6 |
| | | | | Autophagy | 1 | 0.3 | 96.9 |
| | | | | Endocrine function | 1 | 0.3 | 97.1 |
| | | | | Secretion | 1 | 0.3 | 97.4 |
| | | | | Glycoprotein metabolism | 1 | 0.3 | 97.6 |
| | | | | Histone modification | 1 | 0.3 | 97.9 |
| | | | | Metal homeostasis | 1 | 0.3 | 98.2 |
| | | | | Nucleotide metabolism | 1 | 0.3 | 98.4 |
| | | | | Oogenesis | 1 | 0.3 | 98.7 |
| | | | | Protein transport | 1 | 0.3 | 99.0 |
| | | | | Glycoprotein metabolism | 1 | 0.3 | 99.2 |

| | | | | | | | |
|---|---|---|---|---|---|---|---|
| | | | | Steroid metabolism & biosynthesis | 1 | 0.3 | 99.5 |
| | | | | Membrane and substrate transport | 1 | 0.3 | 99.7 |
| | | | | Urea metabolism | 1 | 0.3 | 100.0 |
| **C. Downregulated in 24 h recovery vs. normoxia (144)** | | | | **D. Upregulated in 24 h recovery vs. normoxia (384)** | | | |
| **Functional group** | **# DEGs** | **% of total** | Cum.% | **Functional group** | **# DEGs** | **% of total** | Cum.% |
| Protein synthesis | 18 | 12.5 | 12.5 | Protein synthesis | 39 | 10.2 | 10.2 |
| Immunity | 14 | 9.7 | 22.2 | Immunity | 32 | 8.3 | 18.5 |
| Transposition | 9 | 6.3 | 28.5 | Proteolysis | 25 | 6.5 | 25.0 |
| Adhesion | 7 | 4.9 | 33.3 | Neural function | 20 | 5.2 | 30.2 |
| Proteolysis | 7 | 4.9 | 38.2 | Transcription regulation | 17 | 4.4 | 34.6 |
| Transcription regulation & transcription factors | 7 | 4.9 | 43.1 | Transposition | 17 | 4.4 | 39.1 |
| ECM | 6 | 4.2 | 47.2 | Ciliary function | 16 | 4.2 | 43.2 |
| Cell division & differentiation | 6 | 4.2 | 51.4 | ECM | 15 | 3.9 | 47.1 |
| DNA maintenance | 6 | 4.2 | 55.6 | Signaling | 14 | 3.6 | 50.8 |
| RNA metabolism | 5 | 3.5 | 59.0 | DNA maintenance | 13 | 3.4 | 54.2 |
| Cytoskeleton | 4 | 2.8 | 61.8 | Cell division & differentiation | 11 | 2.9 | 57.0 |
| Ion transport | 4 | 2.8 | 64.6 | Spermatogenesis | 11 | 2.9 | 59.9 |
| Nucleotide metabolism | 4 | 2.8 | 67.4 | PTM | 10 | 2.6 | 62.5 |
| Apoptosis | 3 | 2.1 | 69.4 | Apoptosis | 9 | 2.3 | 64.8 |
| Stress response | 3 | 2.1 | 71.5 | Cytoskeleton | 9 | 2.3 | 67.2 |
| ETS | 3 | 2.1 | 73.6 | ETS | 9 | 2.3 | 69.5 |
| Ciliary activity | 3 | 2.1 | 75.7 | Adhesion | 8 | 2.1 | 71.6 |
| Redox homeostasis | 3 | 2.1 | 77.8 | Detoxification | 8 | 2.1 | 73.7 |
| Chaperone | 2 | 1.4 | 79.2 | Chaperone | 7 | 1.8 | 75.5 |
| Development | 2 | 1.4 | 80.6 | Protein transport | 7 | 1.8 | 77.3 |
| Endocytosis | 2 | 1.4 | 81.9 | Carbohydrate metabolism | 6 | 1.6 | 78.9 |
| Locomotion | 2 | 1.4 | 83.3 | Glycoprotein metabolism | 6 | 1.6 | 80.5 |

| | | | | | | | |
|---|---|---|---|---|---|---|---|
| Protein transport | 2 | 1.4 | 84.7 | Locomotion | 6 | 1.6 | 82.0 |
| Signaling | 2 | 1.4 | 86.1 | Membrane and substrate transport | 6 | 1.6 | 83.6 |
| Oxidoreductase | 2 | 1.4 | 87.5 | Endocytosis | 5 | 1.3 | 84.9 |
| Iron homeostasis | 2 | 1.4 | 88.9 | RNA metabolism | 5 | 1.3 | 86.2 |
| Neural function | 2 | 1.4 | 90.3 | Sulfur metabolism | 5 | 1.3 | 87.5 |
| Peptidoglycan synthesis | 2 | 1.4 | 91.7 | Inflammation | 4 | 1.0 | 88.5 |
| Carbohydrate metabolism | 1 | 0.7 | 92.4 | Ion transport | 4 | 1.0 | 89.6 |
| Detoxification | 1 | 0.7 | 93.1 | Stress response | 4 | 1.0 | 90.6 |
| Endocytosis | 1 | 0.7 | 93.7 | Cofactor metabolism | 3 | 0.8 | 91.4 |
| Glycoprotein metabolism | 1 | 0.7 | 94.4 | Nucleotide metabolism | 3 | 0.8 | 92.2 |
| Histone modification | 1 | 0.7 | 95.1 | Oxidoreductases | 3 | 0.8 | 93.0 |
| Inflammation | 1 | 0.7 | 95.8 | Redox homeostasis | 4 | 1.0 | 94.0 |
| Lipid metabolism | 1 | 0.7 | 96.5 | AA metabolism | 2 | 0.5 | 94.5 |
| mRNA synthesis | 1 | 0.7 | 97.2 | Histone modification | 2 | 0.5 | 95.1 |
| Oogenesis | 1 | 0.7 | 97.9 | Lipid metabolism | 2 | 0.5 | 95.6 |
| Membrane and substrate transport | 1 | 0.7 | 98.6 | Membrane function | 2 | 0.5 | 96.1 |
| Sulfur metabolism | 1 | 0.7 | 99.3 | Oogenesis | 2 | 0.5 | 96.6 |
| Tricarboxylic acid cycle | 1 | 0.7 | 100.0 | Organelle biogenesis | 2 | 0.5 | 97.1 |
| | | | | Secretion | 2 | 0.5 | 97.7 |
| | | | | Tricarboxylic acid cycle | 2 | 0.5 | 98.2 |
| | | | | Angiogenesis | 1 | 0.3 | 98.4 |
| | | | | Ca metabolism | 1 | 0.3 | 98.7 |
| | | | | Cell wall | 1 | 0.3 | 99.0 |
| | | | | Endocrine function | 1 | 0.3 | 99.2 |
| | | | | Nuclear transport | 1 | 0.3 | 99.5 |
| | | | | Sphingolipid metabolism | 1 | 0.3 | 99.7 |
| | | | | Steroid metabolism & biosynthesis | 1 | 0.3 | 100.0 |

**Supplementary Table 5.** Results of the Reactome analysis of the gill transcriptome of *L. capensis*, displaying only significantly up- or downregulated pathways (FDR < 0.1). No significantly enriched pathways were identified in the following comparisons: R24 (24 hours of reoxygenation) vs. normoxia (N), R1 (1 hour of reoxygenation) vs. hypoxia (H), and R24 vs. hypoxia. Additionally, no significantly downregulated pathways were found in the R1 vs N comparison. "# Genes found" refers to the number of identified DEGs, while "# Genes total" refers to the total number of genes in the respective pathway. Rxn – reactions.

| Contrast | Direction | Pathway identifier | Pathway name | #Genes found | #Genes total | P value | FDR | #Rxn found | #Rxn total |
|---|---|---|---|---|---|---|---|---|---|
| **H vs N** | Downregulated | R-HSA-975956 | Nonsense Mediated Decay (NMD) independent of the Exon Junction Complex (EJC) | 60 | 96 | 1.11E-16 | 4.22E-15 | 1 | 1 |
| **H vs N** | Downregulated | R-HSA-72689 | Formation of a pool of free 40S subunits | 63 | 102 | 1.11E-16 | 4.22E-15 | 2 | 2 |
| **H vs N** | Downregulated | R-HSA-192823 | Viral mRNA Translation | 60 | 101 | 1.11E-16 | 4.22E-15 | 2 | 2 |
| **H vs N** | Downregulated | R-HSA-156827 | L13a-mediated translational silencing of Ceruloplasmin expression | 64 | 112 | 1.11E-16 | 4.22E-15 | 3 | 3 |
| **H vs N** | Downregulated | R-HSA-72706 | GTP hydrolysis and joining of the 60S ribosomal subunit | 64 | 113 | 1.11E-16 | 4.22E-15 | 3 | 3 |
| **H vs N** | Downregulated | R-HSA-1799339 | SRP-dependent cotranslational protein targeting to membrane | 60 | 113 | 1.11E-16 | 4.22E-15 | 5 | 5 |
| **H vs N** | Downregulated | R-HSA-72649 | Translation initiation complex formation | 29 | 59 | 1.11E-16 | 4.22E-15 | 2 | 2 |
| **H vs N** | Downregulated | R-HSA-72702 | Ribosomal scanning and start codon recognition | 29 | 59 | 1.11E-16 | 4.22E-15 | 2 | 2 |
| **H vs N** | Downregulated | R-HSA-6791226 | Major pathway of rRNA processing in the nucleolus and cytosol | 71 | 183 | 1.11E-16 | 4.22E-15 | 7 | 7 |
| **H vs N** | Downregulated | R-HSA-927802 | Nonsense-Mediated Decay (NMD) | 61 | 117 | 1.11E-16 | 4.22E-15 | 5 | 6 |
| **H vs N** | Downregulated | R-HSA-72662 | Activation of the mRNA upon binding of the cap-binding complex and eIFs, and subsequent binding to 43S | 29 | 60 | 1.11E-16 | 4.22E-15 | 5 | 6 |
| **H vs N** | Downregulated | R-HSA-156902 | Peptide chain elongation | 60 | 90 | 1.11E-16 | 4.22E-15 | 4 | 5 |
| **H vs N** | Downregulated | R-HSA-975957 | Nonsense Mediated Decay (NMD) enhanced by the Exon Junction Complex (EJC) | 61 | 117 | 1.11E-16 | 4.22E-15 | 4 | 5 |
| **H vs N** | Downregulated | R-HSA-72613 | Eukaryotic Translation Initiation | 64 | 120 | 1.11E-16 | 4.22E-15 | 16 | 21 |
| **H vs N** | Downregulated | R-HSA-8868773 | rRNA processing in the nucleus and cytosol | 73 | 193 | 1.11E-16 | 4.22E-15 | 11 | 15 |

| | | | | | | | | | |
|---|---|---|---|---|---|---|---|---|---|
| **H vs N** | Downregulated | R-HSA-72737 | Cap-dependent Translation Initiation | 64 | 120 | 1.11E-16 | 4.22E-15 | 13 | 18 |
| **H vs N** | Downregulated | R-HSA-156842 | Eukaryotic Translation Elongation | 62 | 95 | 1.11E-16 | 4.22E-15 | 6 | 9 |
| **H vs N** | Downregulated | R-HSA-72312 | rRNA processing | 75 | 203 | 1.11E-16 | 4.22E-15 | 13 | 21 |
| **H vs N** | Downregulated | R-HSA-9711097 | Cellular response to starvation | 63 | 157 | 1.11E-16 | 4.22E-15 | 17 | 28 |
| **H vs N** | Downregulated | R-HSA-72764 | Eukaryotic Translation Termination | 60 | 94 | 1.11E-16 | 4.22E-15 | 3 | 5 |
| **H vs N** | Downregulated | R-HSA-72766 | Translation | 76 | 294 | 1.11E-16 | 4.22E-15 | 54 | 99 |
| **H vs N** | Downregulated | R-HSA-8953854 | Metabolism of RNA | 96 | 729 | 1.11E-16 | 4.22E-15 | 78 | 199 |
| **H vs N** | Downregulated | R-HSA-168273 | Influenza Viral RNA Transcription and Replication | 62 | 152 | 1.11E-16 | 4.22E-15 | 5 | 13 |
| **H vs N** | Downregulated | R-HSA-9735869 | SARS-CoV-1 modulates host translation machinery | 25 | 40 | 1.11E-16 | 4.22E-15 | 1 | 3 |
| **H vs N** | Downregulated | R-HSA-72695 | Formation of the ternary complex, and subsequently, the 43S complex | 28 | 52 | 1.11E-16 | 4.22E-15 | 1 | 3 |
| **H vs N** | Downregulated | R-HSA-9633012 | Response of EIF2AK4 (GCN2) to amino acid deficiency | 61 | 102 | 1.11E-16 | 4.22E-15 | 5 | 16 |
| **H vs N** | Downregulated | R-HSA-2408557 | Selenocysteine synthesis | 60 | 94 | 1.11E-16 | 4.22E-15 | 2 | 7 |
| **H vs N** | Downregulated | R-HSA-9010553 | Regulation of expression of SLITs and ROBOs | 64 | 172 | 1.11E-16 | 4.22E-15 | 5 | 19 |
| **H vs N** | Downregulated | R-HSA-2408522 | Selenoamino acid metabolism | 64 | 118 | 1.11E-16 | 4.22E-15 | 5 | 23 |
| **H vs N** | Downregulated | R-HSA-168255 | Influenza Infection | 63 | 172 | 1.11E-16 | 4.22E-15 | 11 | 58 |
| **H vs N** | Downregulated | R-HSA-376176 | Signaling by ROBO receptors | 65 | 218 | 1.11E-16 | 4.22E-15 | 7 | 59 |
| **H vs N** | Downregulated | R-HSA-71291 | Metabolism of amino acids and derivatives | 78 | 376 | 1.11E-16 | 4.22E-15 | 22 | 248 |
| **H vs N** | Downregulated | R-HSA-9754678 | SARS-CoV-2 modulates host translation machinery | 25 | 53 | 3.33E-16 | 1.23E-14 | 1 | 6 |
| **H vs N** | Downregulated | R-HSA-422475 | Axon guidance | 71 | 558 | 6.15E-12 | 2.21E-10 | 26 | 297 |
| **H vs N** | Downregulated | R-HSA-2262752 | Cellular responses to stress | 86 | 779 | 4.41E-11 | 1.54E-09 | 93 | 486 |

| | | | | | | | | | |
|---|---|---|---|---|---|---|---|---|---|
| **H vs N** | Downregulated | R-HSA-9675108 | Nervous system development | 71 | 584 | 4.60E-11 | 1.56E-09 | 26 | 323 |
| **H vs N** | Downregulated | R-HSA-9692914 | SARS-CoV-1-host interactions | 27 | 109 | 5.55E-11 | 1.83E-09 | 6 | 48 |
| **H vs N** | Downregulated | R-HSA-8953897 | Cellular responses to stimuli | 86 | 793 | 1.06E-10 | 3.40E-09 | 93 | 517 |
| **H vs N** | Downregulated | R-HSA-9678108 | SARS-CoV-1 Infection | 30 | 154 | 1.33E-09 | 4.11E-08 | 12 | 156 |
| **H vs N** | Downregulated | R-HSA-9705683 | SARS-CoV-2-host interactions | 32 | 221 | 3.29E-07 | 9.87E-06 | 10 | 67 |
| **H vs N** | Downregulated | R-HSA-1430728 | Metabolism | 160 | 2150 | 7.06E-07 | 2.12E-05 | 237 | 2044 |
| **H vs N** | Downregulated | R-HSA-9694516 | SARS-CoV-2 Infection | 36 | 316 | 1.35E-05 | 3.92E-04 | 17 | 208 |
| **H vs N** | Downregulated | R-HSA-9824446 | Viral Infection Pathways | 79 | 980 | 6.95E-05 | 0.00194584 | 89 | 644 |
| **H vs N** | Downregulated | R-HSA-6790901 | rRNA modification in the nucleus and cytosol | 11 | 60 | 3.74E-04 | 0.01046335 | 4 | 8 |
| **H vs N** | Downregulated | R-HSA-163200 | Respiratory electron transport, ATP synthesis by chemiosmotic coupling, and heat production by uncoupling proteins. | 17 | 127 | 4.50E-04 | 0.0121625 | 17 | 28 |
| **H vs N** | Downregulated | R-HSA-1253288 | Downregulation of ERBB4 signaling | 4 | 11 | 0.00277726 | 0.07220863 | 5 | 5 |
| **H vs N** | Downregulated | R-HSA-611105 | Respiratory electron transport | 13 | 103 | 0.00332034 | 0.08358227 | 14 | 19 |
| **H vs N** | Downregulated | R-HSA-6799198 | Complex I biogenesis | 9 | 57 | 0.00334329 | 0.08358227 | 11 | 13 |
| **H vs N** | Upregulated | R-HSA-1428517 | The citric acid (TCA) cycle and respiratory electron transport | 13 | 238 | 6.33E-05 | 0.06294941 | 24 | 67 |
| **R1 vs N** | Upregulated | R-HSA-5675482 | Regulation of necroptotic cell death | 6 | 39 | 4.72E-05 | 0.0465962 | 9 | 21 |
| **R1 vs N** | Upregulated | R-HSA-5213460 | RIPK1-mediated regulated necrosis | 6 | 45 | 1.03E-04 | 0.0507073 | 10 | 29 |
| **R1 vs N** | Upregulated | R-HSA-168927 | TICAM1, RIP1-mediated IKK complex recruitment | 4 | 19 | 2.79E-04 | 0.09183038 | 1 | 3 |

**Supplementary Table 6.** Results of the Reactome analysis of the digestive gland transcriptome of *L. capensis*, displaying only significantly up- or downregulated pathways (FDR < 0.1). No significantly downregulated pathways were identified in the following comparisons: hypoxia (H) vs. normoxia (N), R24 (24 hours of reoxygenation) vs. N. No significantly downregulated pathways were identified in the following comparisons: R24 v. N and R24 vs. A. "# Genes found" refers to the number of identified DEGs, while "# Genes total" refers to the total number of genes in the respective pathway. Rxn – reactions.

| Contrast | Direction | Pathway identifier | Pathway name | #Genes found | #Genes total | P value | FDR | #Rxn found | #Rxn total |
|---|---|---|---|---|---|---|---|---|---|
| **H vs N** | Upregulated | R-HSA-8851708 | Signaling by FGFR2 IIIa TM | 5 | 23 | 2.62E-04 | 0.0965289 | 2 | 2 |
| **H vs N** | Upregulated | R-HSA-1679131 | Trafficking and processing of endosomal TLR | 4 | 13 | 3.01E-04 | 0.0965289 | 2 | 7 |
| **H vs N** | Upregulated | R-HSA-1839126 | FGFR2 mutant receptor activation | 6 | 38 | 3.52E-04 | 0.0965289 | 17 | 18 |

**Supplementary Table 7.** Significantly enriched pathways identified through MetaboAnalyst analysis of *L. capensis* gill metabolome profiles under normoxic conditions versus 1 hour of reoxygenation. Only pathways with a False Discovery Rate (FDR) < 0.1 are included. "Total Compounds" refers to the total number of compounds associated with each pathway, while "Hits" indicates the number of metabolites detected in *L. capensis* gills that are linked to each pathway.

| **Pathway** | **Total compounds** | **Hits** | **Raw p** | **FDR** | **Impact** |
|---|---|---|---|---|---|
| Citrate cycle (TCA cycle) | 20 | 4 | 0.00062 | 0.0096 | 0.18 |
| Butanoate metabolism | 14 | 4 | 0.00876 | 0.0761 | 0.2 |
| Pyruvate metabolism | 23 | 3 | 0.01187 | 0.0761 | 0.18 |
| Cysteine and methionine metabolism | 32 | 5 | 0.01227 | 0.0761 | 0.21 |

**Supplementary Figure 1.** GO (A, B) and KEGG (C,D) pathway enrichment in the gills of *L. capensis* after 1 h of reoxygenation relative to the hypoxic state. Color indicates the significance ($p_{adj}$) and size of the symbols – the number of DEGs found in the respective pathway. X axis shows Gene Ratio for each pathway, depicting the ratio of differentially expressed genes to all genes for this GO term or KEGG pathway.

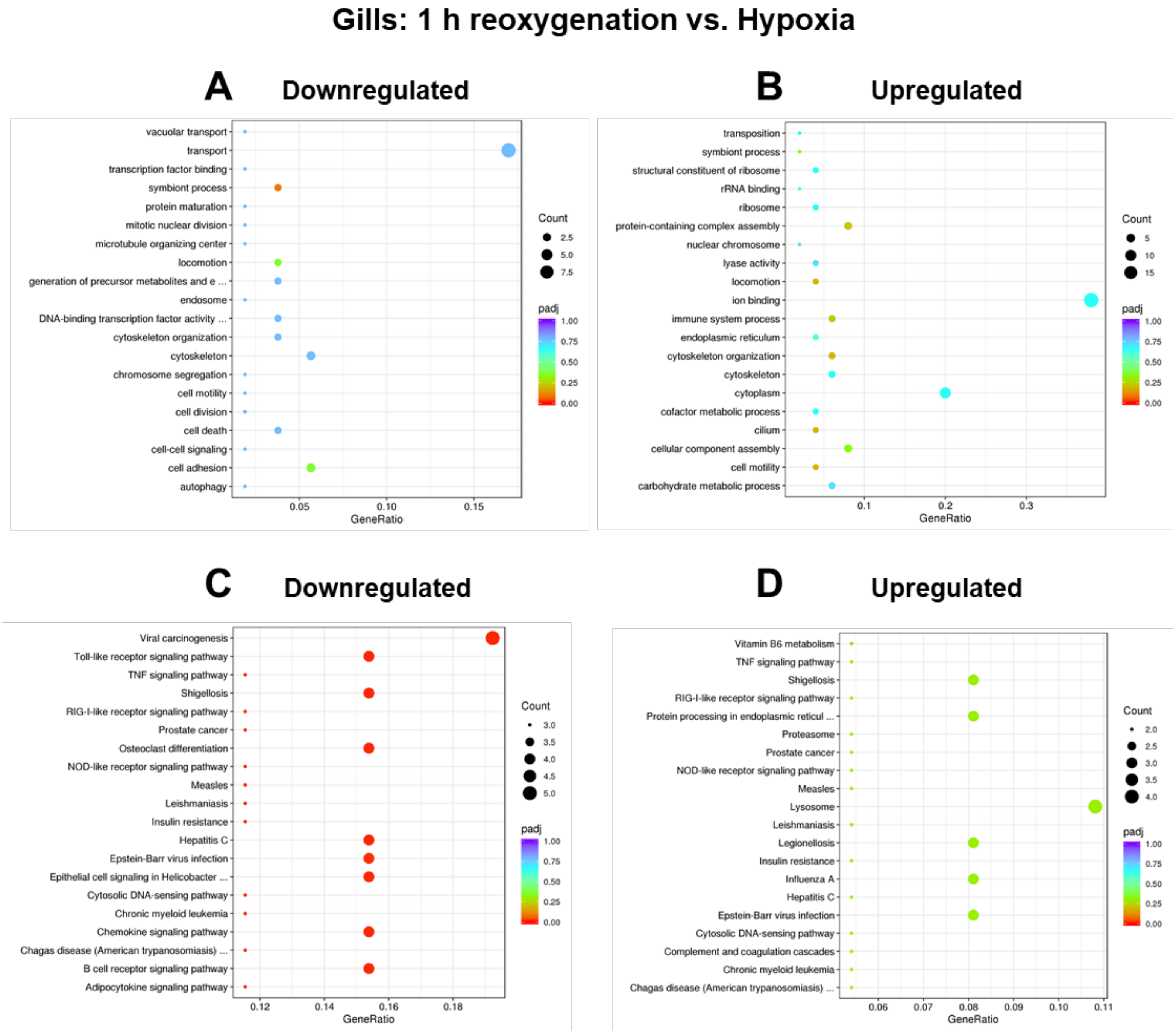

**Supplementary Figure 2.** GO (A, B) and KEGG (C,D) pathway enrichment in the gills of *L. capensis* after 24 h of reoxygenation relative to the hypoxic state. Color indicates the significance ($p_{adj}$) and size of the symbols – the number of DEGs found in the respective pathway. X axis shows Gene Ratio for each pathway, depicting the ratio of differentially expressed genes to all genes for this GO term or KEGG pathway.

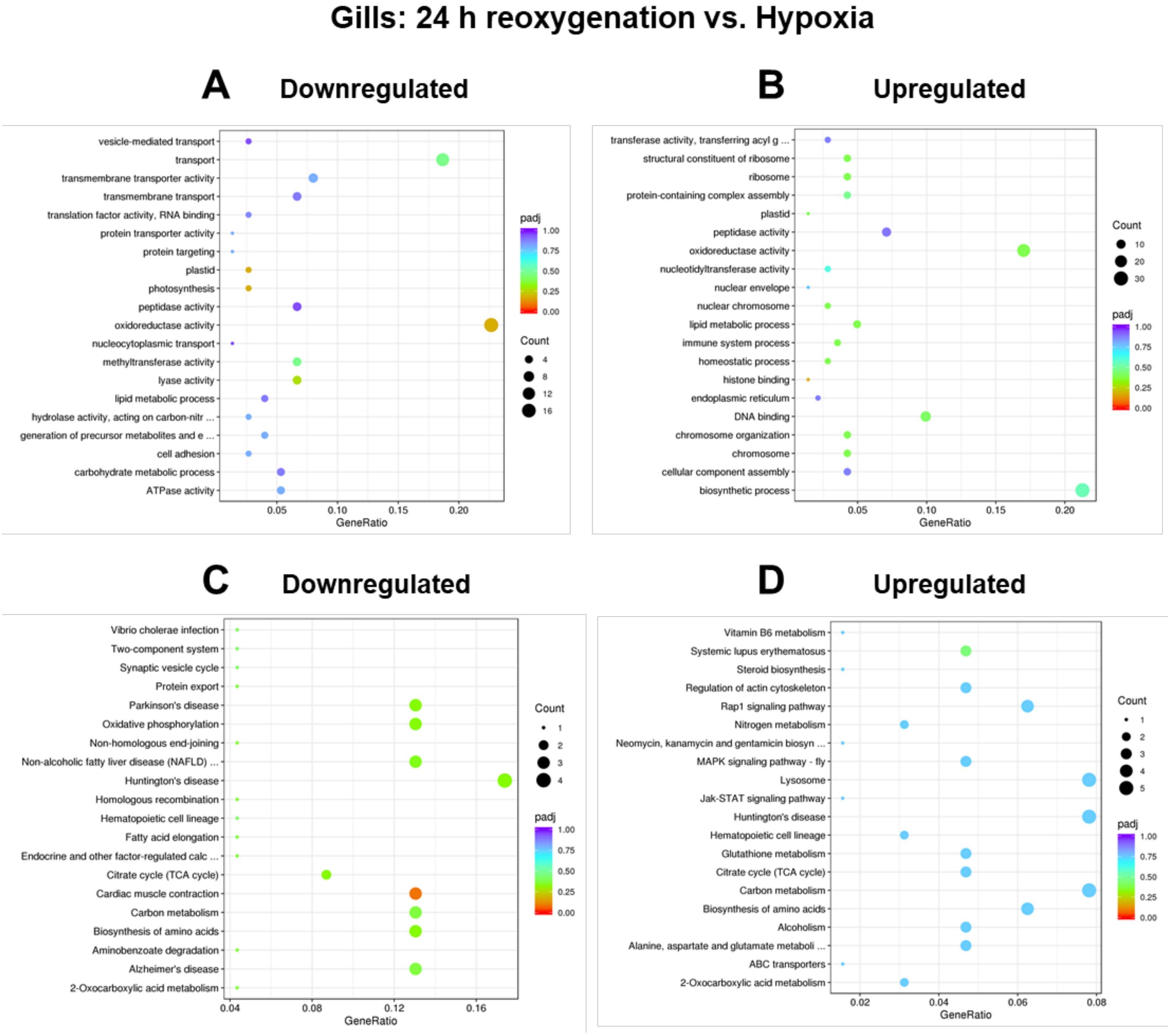

**Supplementary Figure 3.** GO (A, B) and KEGG (C,D) pathway enrichment in the hypoxic digestive gland of *L. capensis* relative to the normoxic control. Color indicates the significance ($p_{adj}$) and size of the symbols – the number of DEGs found in the respective pathway. X axis shows Gene Ratio for each pathway, depicting the ratio of differentially expressed genes to all genes for this GO term or KEGG pathway.

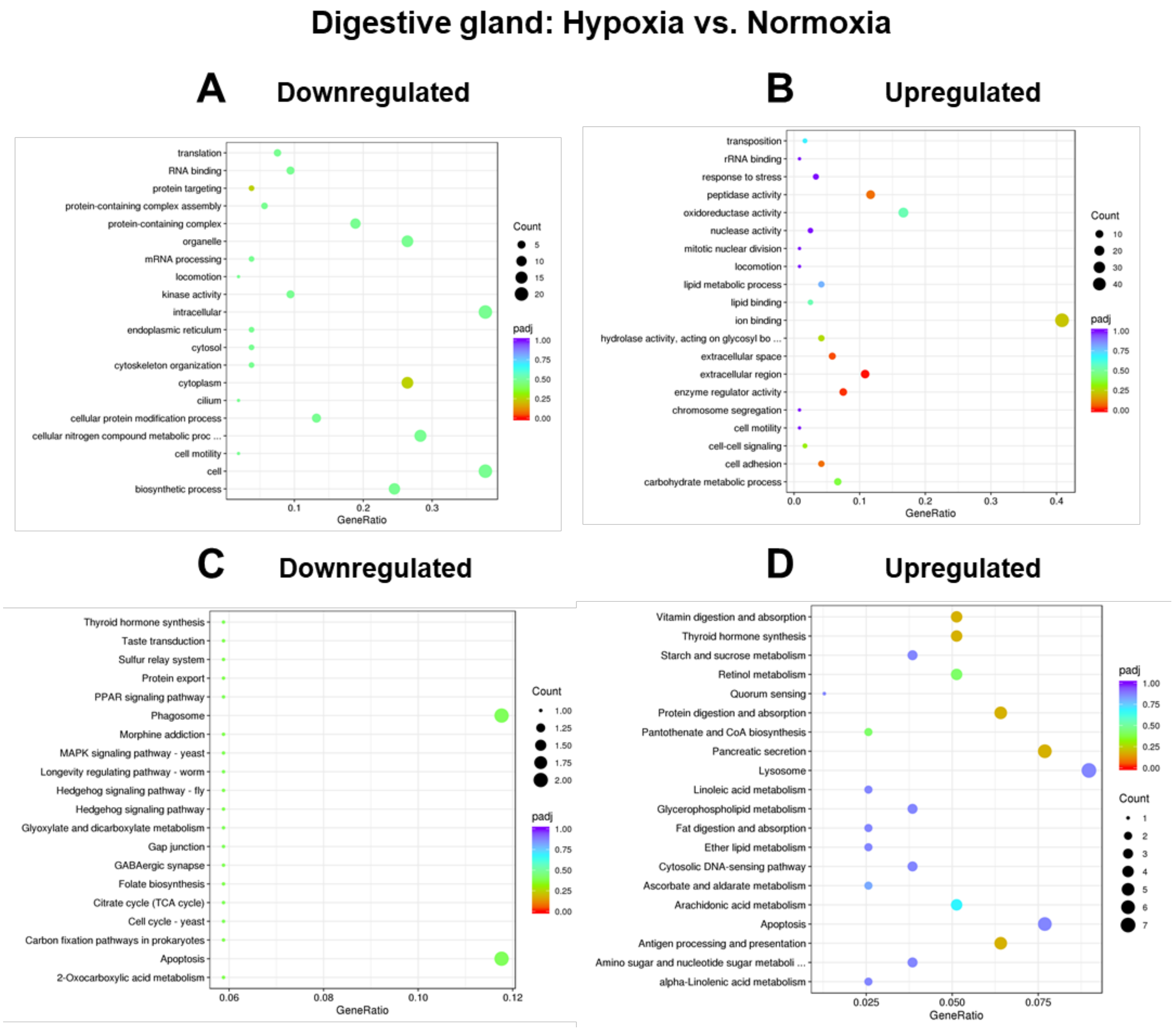

**Supplementary Figure 4.** GO (A, B) and KEGG (C,D) pathway enrichment in the digestive gland of *L. capensis* after 24 h of reoxygenation relative to the normoxic control. Color indicates the significance ($p_{adj}$) and size of the symbols – the number of DEGs found in the respective pathway. X axis shows Gene Ratio for each pathway, depicting the ratio of differentially expressed genes to all genes for this GO term or KEGG pathway.

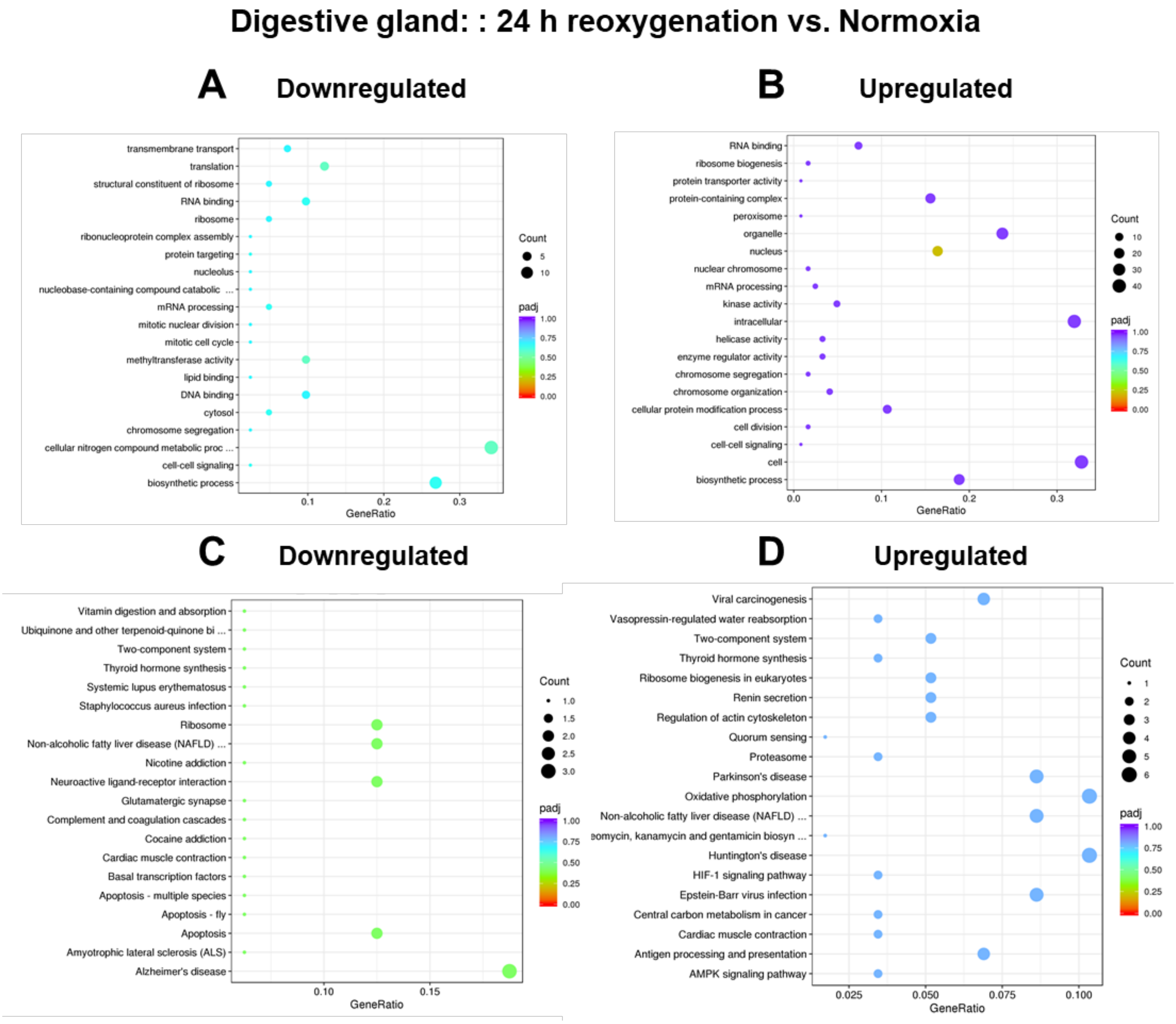

**Supplementary Figure 5.** GO (A, B) and KEGG (C,D) pathway enrichment in the digestive gland of *L. capensis* after 24 h of reoxygenation relative to the hypoxic state. Color indicates the significance ($p_{adj}$) and size of the symbols – the number of DEGs found in the respective pathway. X axis shows Gene Ratio for each pathway, depicting the ratio of differentially expressed genes to all genes for this GO term or KEGG pathway.

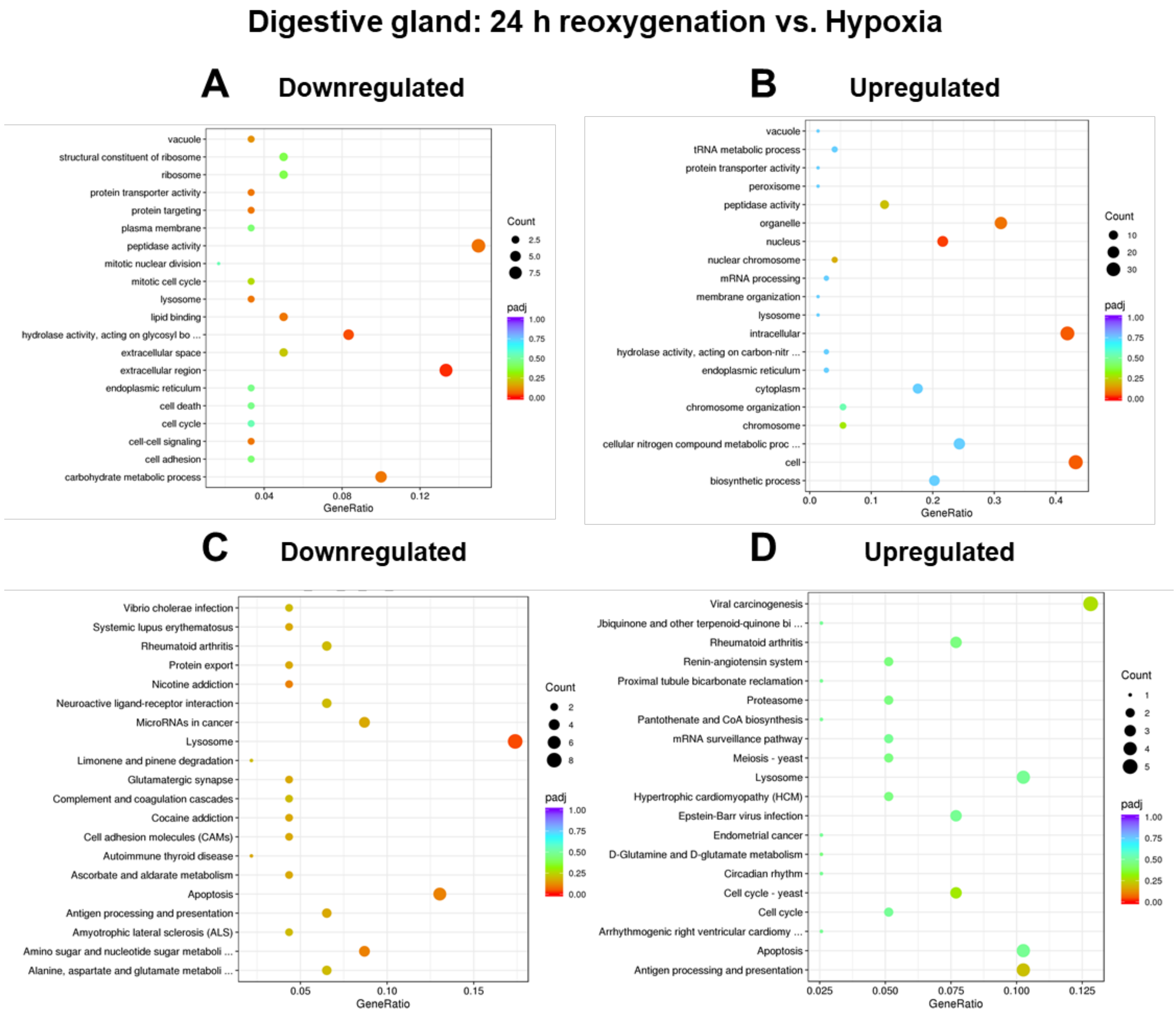
